# A scoping review of evidence of naturally occurring Japanese encephalitis infection in vertebrate animals other than humans, ardeid birds and pigs

**DOI:** 10.1101/2024.04.18.589998

**Authors:** Zoë A. Levesque, Michael Walsh, Cameron Webb, Ruth Zadoks, Victoria J. Brookes

## Abstract

Japanese encephalitis virus (JEV) is the leading cause of human encephalitis in Asia. JEV is a vector-borne disease, mainly transmitted by *Culex* mosquitoes, with *Ardeidae* birds as maintenance hosts and pigs as amplifying hosts. Other vertebrate animal hosts have been suggested to play a role in the epidemiology of JEV. This scoping review followed PRISMA guidelines to identify species in which evidence of naturally occurring JEV infection was detected in vertebrates other than ardeid birds, pigs and people. Following systematic searches, 4372 records were screened, and data were extracted from 62 eligible studies. Direct evidence (virus, viral antigen or viral RNA) of JEV infection was identified in a variety of mammals and birds (not always identified to the species level), including bats, passerine birds (family Turdidae), livestock (cattle [*Bos taurus*] and a goat [*Capra hircus*]), carnivores (two meerkats [*Suricata suricatta*]), and one horse (*Equus caballus*). Bat families included Pteropodidae, Vespertilionidae, Rhinolophidae, Miniopteridae, Hipposideridae. Indirect evidence suggestive of JEV infection (antibodies) was detected in several mammalian and avian orders, as well as reported in two reptile species. However, a major limitation of the evidence of JEV infection identified in this review was diagnostic test accuracy, particularly for serological testing. Studies generally did not report diagnostic sensitivity or specificity. Although we hypothesise that bats and passerine birds could play an underappreciated role in JEV epidemiology, development of diagnostic tests to differentiate JEV from other orthoflaviviruses will be critical for effective surveillance in these, as well as the companion and livestock species that could be used to evaluate JEV control measures in currently endemic regions.

## 1. INTRODUCTION

Japanese encephalitis (JE) is caused by the zoonotic, mosquito-borne Japanese encephalitis virus (JEV), a single stranded RNA virus of the genus *Orthoflavivirus* [1–3]. Japanese encephalitis was first described in Japan in 1924 and is now endemic in >20 countries in South-East Asia and the Western Pacific region [4]. Although JE incidence is uncertain due to limited surveillance in many countries in which JEV circulates, JEV is considered to be the leading cause of viral encephalitis in Asia, with an estimated 67,900—100,000 JE cases annually, of which 75% occur in children <14 years old [4, 5]. Manifestations of JEV infection vary from asymptomatic (approximately 98% of infections) or nonspecific febrile illness to severe meningoencephalitis, acute flaccid paralysis, and death [1]. Among those with reported symptomatic infection, JE is estimated to have a case fatality rate of 20—30%, with up to 50% of survivors reporting long term neurological sequelae including ‘locked-in syndrome’ in which a JE survivor is cognitively aware but paralysed [6, 7].

Japanese encephalitis virus is transmitted primarily by mosquitoes of genus *Culex* [8], and waterbirds of the family *Ardeidae* are the major maintenance hosts in endemic transmission cycles [9]. In pigs, JEV infection also causes a viraemia sufficient to infect mosquitoes for ongoing transmission. Pigs are considered amplifying hosts and major contributors to the local transmission risk of JE to people in endemic regions with high densities of domestic pigs [10, 11]. Infection in pigs can be asymptomatic or manifest as reproductive failure characterised by abortions, stillbirths, and mummified foetuses [10,11]; for example, stillbirths and abortions affected up to 50% of unvaccinated pregnant sows during a JEV outbreak in Japan [12]. People are dead-end hosts because their associated viremia is not sufficient for transmission to mosquitoes [14].

Japanese encephalitis virus and other mosquito-borne orthoflaviviruses, such as Zika virus, dengue haemorrhagic fever virus, and St Louis (SLEV), West Nile (WNV) and Murray Valley encephalitis viruses, are ongoing global health threats to human and animal health due to their mutability and propensity for emergence and re-emergence [12]. There is growing concern about the possibility of emergence of JEV globally [13–15]. The mechanisms of emergence are not well understood. Climate change and associated changes in temperature and rainfall, and increasingly frequent extreme weather events are all likely to influence the abundance and distribution of mosquito vectors [16, 17]. Changes in the distribution of JEV circulation can also be influenced by factors such as bird migration, movement of infected domesticated hosts, and land use changes [18]. For example, an increase in urban pig farming has resulted in increased JEV circulation in urban and peri-urban settings [19], and expansion of rice farming has resulted in more standing water, thus increasing vector presence [20].

Emergence and re-emergence of JEV might also be influenced by the availability of recognized or alternative vectors and hosts. For example, whilst *Culex tritaeniorhynchus* is the predominant vector mosquito across Southeast Asia, other *Culex* species are regionally predominant or suggested as secondary vectors, including *Culex vishnui, Culex annulirostris*, *Culex palpalis*, *Culex quinquefasciatus* and *Culex sitiens*, and JEV has also been isolated from other mosquito genera such as *Aedes, Anopheles,* and *Mansonia* [21]. In the recent geographic expansion of JEV across eastern and southern Australia, where *Cx. tritaeniorhynchus* has not been detected (unlike in northern Australia [22]), the major JEV vector was considered to be *Culex annulirostris* [23].

The occurrence of JE in people in regions without high densities of ardeid birds or pigs also suggests the presence of alternative maintenance and amplifying hosts (competent hosts) [4, 24, 25]. For example, epidemiological investigations in India indicate an important role of domestic chickens [19]. The role of domestic poultry (orders Galliformes and Anseriformes) as amplifying hosts has been suggested elsewhere too but is poorly understood due to varying levels of reported seroprevalence and uncertainty about levels of viraemia [26]. Although most studies of JEV epidemiology focus on competent hosts that contribute to virus transmission [27], there could also be an epidemiologically important role of non-competent hosts, i.e. dead-end hosts that become infected but develop insufficient viraemia for ongoing transmission. These species might contribute to a dilution effect by reducing viral circulation among hosts within maintenance communities and their mosquito vectors, as well as have a role in JEV surveillance. Examples include domestic animals such as dogs, horses and cattle, as well as wild bird or mammal species [28–30].

The objective of the current study was to systematically review and collate evidence of naturally occurring JEV infection in vertebrate species other than humans, ardeid birds and pigs. The aim of the review is to inform surveillance as well as further research to understand the potential contribution of alternative host species in JEV epidemiology and JE mitigation strategies.

## 2. MATERIALS AND METHODS

### 2.1. Study overview

A PRISMA-guided scoping review [31] was conducted to identify evidence of naturally occurring JEV infection in vertebrate animals other than humans, ardeid birds (herons, egrets and bitterns) and pigs, either by direct detection (virus, viral antigen, or viral RNA) suggesting current infection, or indirect detection (antibody) indicating previous exposure.

### 2.2. Search strategy

Search terms included: “Japanese encephalitis virus” OR JEV OR JE, AND detection OR detected (details in Table S1). Searches for eligible records were conducted in April 2023 and January 2024 in Web of Science, Scopus, and ProQuest Central databases. The first 100 search results from Google Scholar were also screened with a date range of 1935-1980 to identify records published without inclusion in the searched databases. To validate and supplement the database search strategy, records were also identified by snowballing, starting with the bibliographies of two recent reviews on the pathogenesis of JEV infection [33,34].

### 2.3. Eligibility criteria

Records were eligible if they were peer-reviewed and described primary research in which evidence of naturally occurring JEV infection was found in any vertebrate animal other than humans, ardeid birds or pigs using a laboratory test. Animals could be sampled for any reason, including population-level surveillance and outbreak investigation, or diagnosis of disease in individual animals. Investigations of vaccinated animals (for example, horses) were excluded unless studies were investigating natural infection. Laboratory tests included methods to detect virus, viral antigen or viral RNA or JEV antibody. All observational study designs were eligible, including case reports, but experimental studies in which animals were inoculated with JEV were excluded. Records published in English in any year from any location were eligible.

### 2.4. Screening and data charting

Following removal of duplicates, two reviewers (ZL and VB) screened records for eligibility based on title and abstract (Level 1) then full manuscript text (Level 2).

Following screening, data were extracted from eligible records (Level 3) including: month and year of sampling, country and region of study, species investigated, number of animals tested (including number positive), the animals’ circumstances (owned, captive, farmed, wild, and whether sampling had been targeted to identify infection in animals with clinical signs of disease), and the method of detection of JEV infection. Data relating to pigs and ardeid birds were also charted if they had been included in eligible studies to provide the full range and context of the animals investigated in the studies.

### 2.5. Analysis

Data were summarised narratively, and descriptive statistical analyses were conducted in R [32] with packages plyr [33], ggplot2 [34], meta [35], and metafor [36]. Forest plots were used to visualise point estimates of JEV prevalence (direct detection) and JEV seroprevalence (antibody detection) with 95% confidence intervals (Clopper-Pearson method) in animals grouped by the lowest taxonomic rank that was feasible given the aggregation of species reported in the studies. Estimates were not combined across studies due to the methodological heterogeneity of studies (for example, different sampling strategies, and different methods used for detection of infection). A probability density plot was used to visualise detection method usage throughout the period during which studies were conducted.

## 3. RESULTS

### 3.1. Search outcomes

The search strategy identified 4,359 records (Figure 1). Following removal of 1,534 duplicates, 2,748 and 28 records were removed during Level 1 and 2 screening, respectively. Most excluded records were not relevant to the review (did not investigate evidence for naturally occurring JEV infection in vertebrate animals other than people, ardeid birds and pigs). From the records identified by snowballing (n = 13), six were excluded at Level 1, and seven were eligible for data extraction. A total of 62 records were eligible for data extraction (extracted data are available in Table S2).

**Figure 1.**
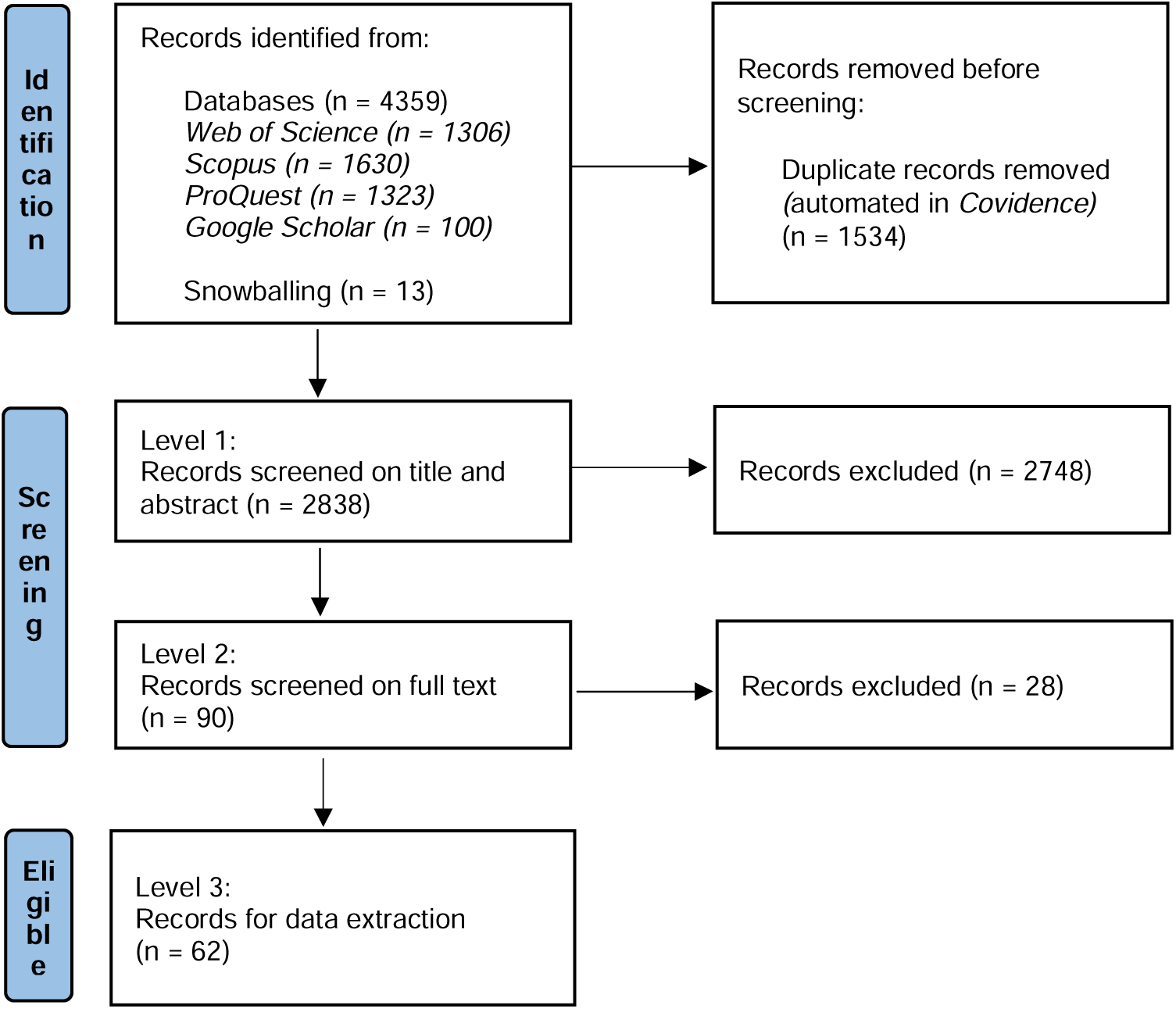
PRISMA diagram of records at each level in a scoping review of direct and indirect evidence of naturally occurring Japanese encephalitis virus infection in vertebrate animals other than humans, ardeid birds and pigs.

### 3.2. Study distribution

Studies were published from 1947 [37] to 2023 [38] and were mainly conducted in Asia (n = 55; Figure S1), with the highest number in Japan (n = 17), followed by India (n = 11), and China (n = 8). Three studies were conducted in locations outside the current known distribution of JEV circulation: North America (n = 1; United States [39]) and Europe (n = 2; Italy [40, 41]). Data were collected over periods of 1—6 years from 1946—2020 inclusive; the distribution of the earliest year of data collection in each study (for studies in which this was recorded [n = 57]) or the date of publication by country is shown in Figure 2. Naturally occurring JEV infection in vertebrate animals other than humans, ardeid birds and pigs was first investigated in Japan in the 1940s, followed by Malaysia in the 1950s. Since then, the number of studies increased each decade, peaking in 2000-2010, then declining in 2011— 2020.

**Figure 2.**
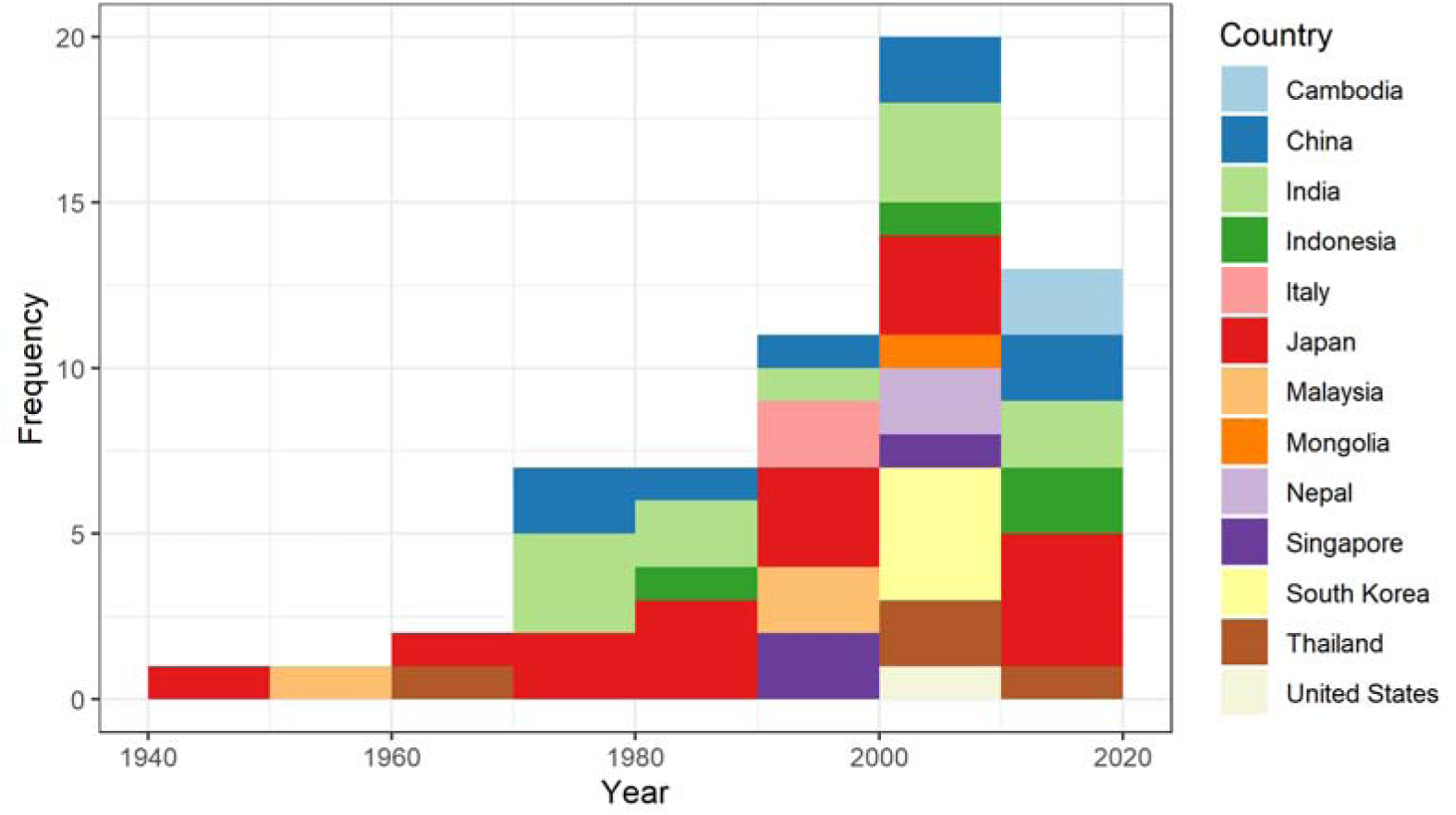
Histogram of the starting year of data collection (or date of publication for 5 studies) by country in a scoping review of direct and indirect evidence of naturally occurring Japanese encephalitis virus infection in vertebrate animals other than humans, ardeid birds and pigs.

### 3.3. Laboratory tests

Tests for direct and indirect detection of JEV infection were conducted in 29% (n = 18) and 85% of studies (n = 51), respectively. The methods used varied according to the years in which the study was conducted (Figure 3).

**Figure 3.**
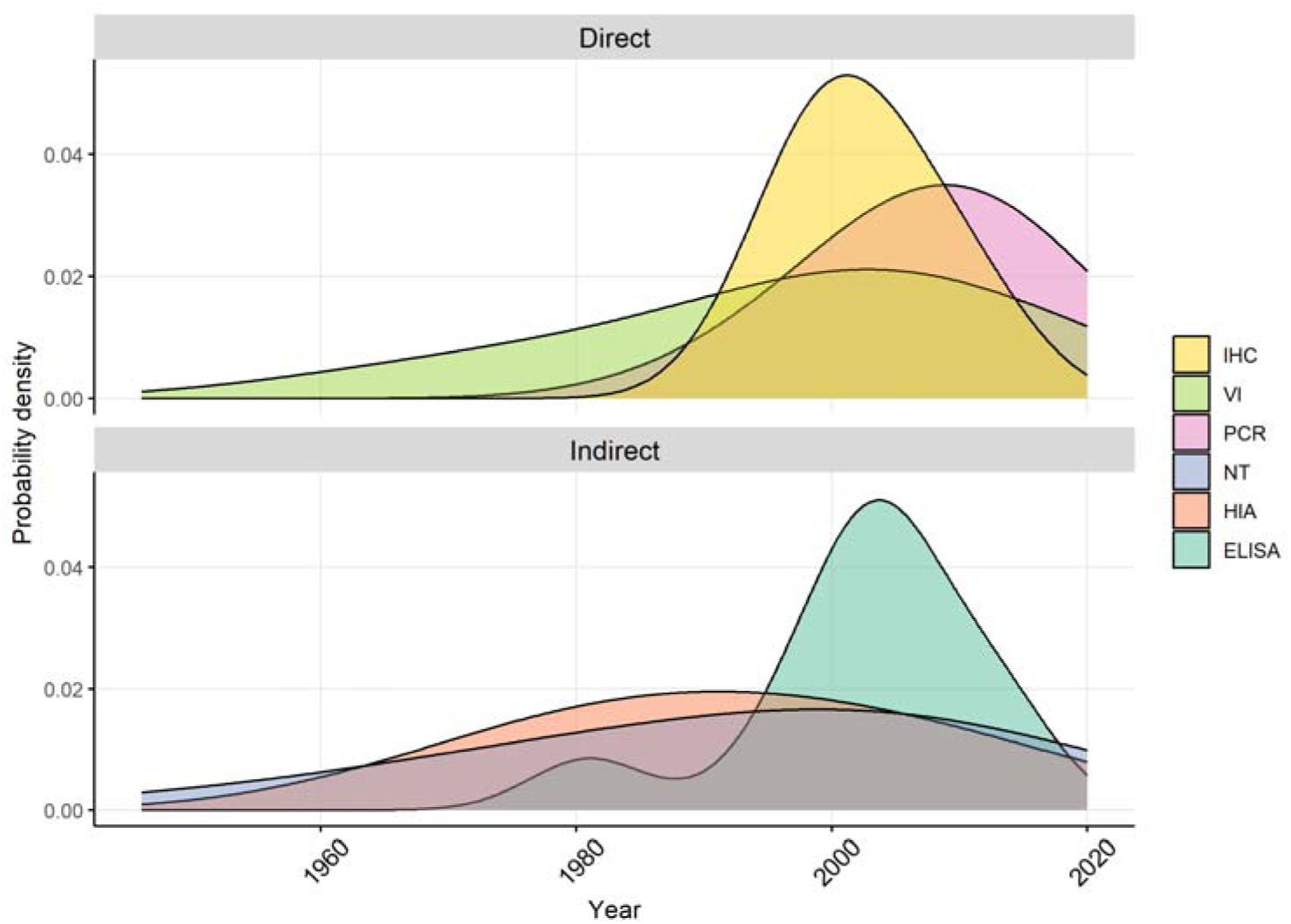
Density plots of methods used in studies identified in a scoping review of direct and indirect evidence of naturally occurring Japanese encephalitis virus infection in vertebrate animals other than humans, ardeid birds and pigs. IHC = immunohistochemistry, VI = virus isolation, PCR = polymerase chain reaction, NT = neutralisation test, HIA = haemagglutinin inhibition test, ELISA = enzyme-linked immunosorbent assay

Direct methods of detection of JEV infection included virus isolation which was conducted in five studies (using inoculation of live mice or Vero, MK2 and *Ae. albopictus* C6/36 cell cultures) with subsequent phylogenetic analyses or transmission confirmation studies [42–45]. Immunohistochemistry (IHC) was conducted in three studies to detect the presence of JEV antigen in tissues from animals that had exhibited neurological signs [42, 43, 46]. Initial detection by IHC was followed by visualisation with electron microscopy in two deceased meerkats (*Suricata suricatta*) in Thailand [46]. Immunohistochemistry was also used to detect virus in archived samples from passerine birds from Tuscany, Italy, following its detection in mosquitoes in the same region in 1997-2000 [40, 41]. Reverse transcriptase-PCR (RT-PCR) tests were conducted from the 1990s onwards in 17 studies. Reported primers targeted the non-structural (NS) protein1, NS3, NS5 and E (envelope) gene regions of JEV. Indirect tests included haemagglutination-inhibition assays (HIA; n = 26 studies), virus neutralisation tests (NT; n = 38 studies), enzyme-linked immunosorbent assays (ELISA; n = 10 studies), immunofluorescence assay (IF, n = 1), and immunochemical staining (n = 1). Studies in which HIA and NT tests were conducted used a range of JEV strains including, most commonly, the Nakayama strain (n = 15), followed by P20778 (n = 5) and JaGAr01 (n = 3). ELISAs were designed to detect antibody rather than virus and used envelope protein antigen (when reported) from a range of JEV strains (including Beijing 01 and SA14) as well as vaccine strains (Bundo, Cui, Konishi, Shimoda; further information in Table S2). The diagnostic performance of tests or testing strategies was reported in only two studies. In a study from Japan, authors reported the diagnostic sensitivity and specificity of an ELISA as 82% and 96%, respectively, when compared to a virus neutralisation test as a gold standard [47]. In another study from Japan, a testing strategy (diagnostic algorithm using ELISA, IF, and NT) was used in which tests were conducted sequentially and dependent on the previous test result in the algorithm, and authors reported 100% accuracy in differentiating detection of WNV exposure from JEV exposure [48]. Throughout most of the studies, however, authors recognised the limitations of potentially poor test performance due to cross-reactivity with other flaviviruses, especially when using antibody tests. Serial testing was often conducted to increase specificity (for example, authors sometimes stated that more specific NT tests were used to confirm HIA positive samples [NT was use sequentially in 62% of studies using HIA]). Due to the overall lack of reported diagnostic sensitivity and specificity in studies in this review, we make no assumptions about test performance and report apparent prevalence (direct testing) and apparent seroprevalence (indirect testing).

### 3.4. Evidence of JEV infection

Reported direct and indirect evidence is described below according to the finest-resolved animal taxa available in the studies (class and order or family; Figures 4—9, and S2—3), and is shown by country in the Supplementary Material (Figures S4—6).

**Figure 4.**
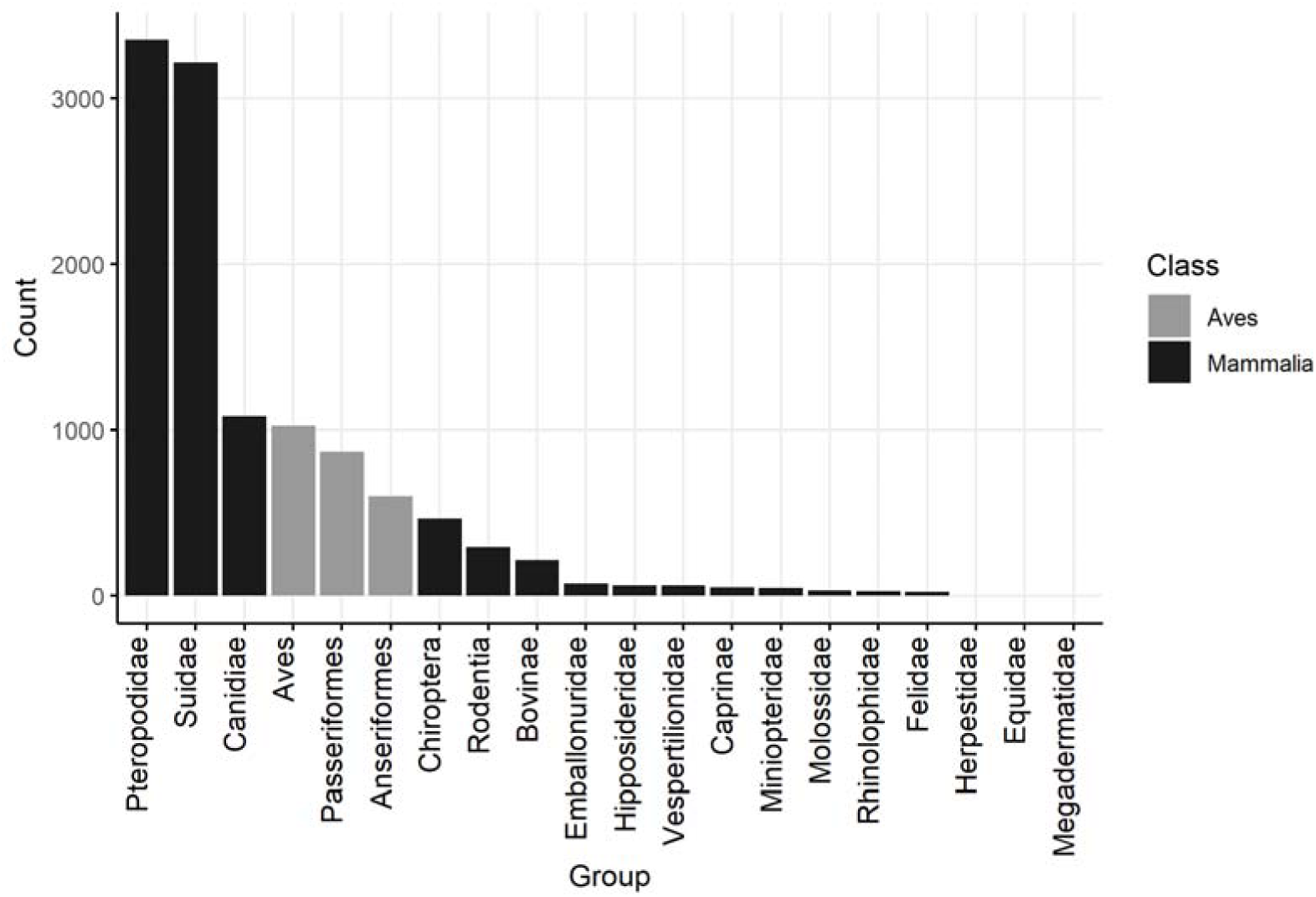
Barplots of the number of animals tested for direct evidence of naturally occurring JEV infection in studies of vertebrate animals other than humans, ardeid birds and pigs. Suidae are included from studies in which vertebrate animals other than humans, ardeid birds or pigs were also tested.

#### 3.4.1. Direct evidence of JEV

In the studies identified, 9004 mammalian (including 3215 Suidae) and 2491 avian (no Ardeidae specified) individual animals were tested for direct evidence of JEV infection (virus, viral antigen or viral RNA). Reptiles, fish and amphibia were not tested. The largest family of mammals tested was Pteropodidae (pteropodid bats, also known as Megachiroptera or megabats, fruit bats, old world bats, and flying foxes; n = 3346) (Figures 4 and 5). Direct evidence of JEV infection was detected in pteropodid bats in Indonesia and India but not in Thailand or China. In a survey from 11 provinces in Indonesia, JEV viral RNA was detected by RT-PCR targeting the NS3 gene in blood samples from 61 of 2805 live, wild-caught pteropodid bats (2.3%) including the genera *Cynopterus*, *Dobsonia*, *Eonycteris*, *Macroglossus*, *Pteropus*, *Rousettus*, and *Thoopterus* [49]. Viral RNA was also detected in pteropodid bats in an earlier study in Indonesia (up to 43% of *Balionycterus maculata*; other species were up to 9% prevalence) [50, 51]. The first detection of JEV infection in bats in India was in 2020, associated with a mortality event in which 52 pteropodid bats were found dead in Gorakhpur district, Uttar Pradesh, a province which has a high incidence of JE in people and JEV circulating in domestic pigs [52, 53]. Brain tissue samples from 8 bats were tested by conventional RT-PCR and TaqMan real-time RT-PCR using primers targeting the JEV envelope and NS1 genes. Five bats were positive by TaqMan real-time RT-PCR (62%, with 8—18 copies/reaction). Other bat families in which evidence of JEV infection was detected included Vespertilionidae, Rhinolophidae, Miniopteridae, and Hipposideridae in Indonesia and China [49, 54]. Complete genome sequencing was conducted on eight different isolates from bats (Rhinolophidae, Miniopteridae, and Vespertilionidae, as well as Pteropodidae) which were sampled in China in Yunnan, Hainan, Guangdong, and Hunan provinces from 1986—2007 [54, 55]. The sequence data were consistent with JEV Genotype III, showed 99.4—99.9 % genetic homogeneity in the full-length nucleotide sequences, and were phylogenetically clustered into the same subgroup despite wide spatial and temporal sampling. JEV was not detected in bat families Emballonuridae (Indonesia and China), Megadermatidae (Indonesia), and Molossidae (Indonesia; Figure 6) [49, 56]. In all the studies of bats identified above except Dhanze et al. (2022), sampled bats had appeared healthy.

**Figure 5.**
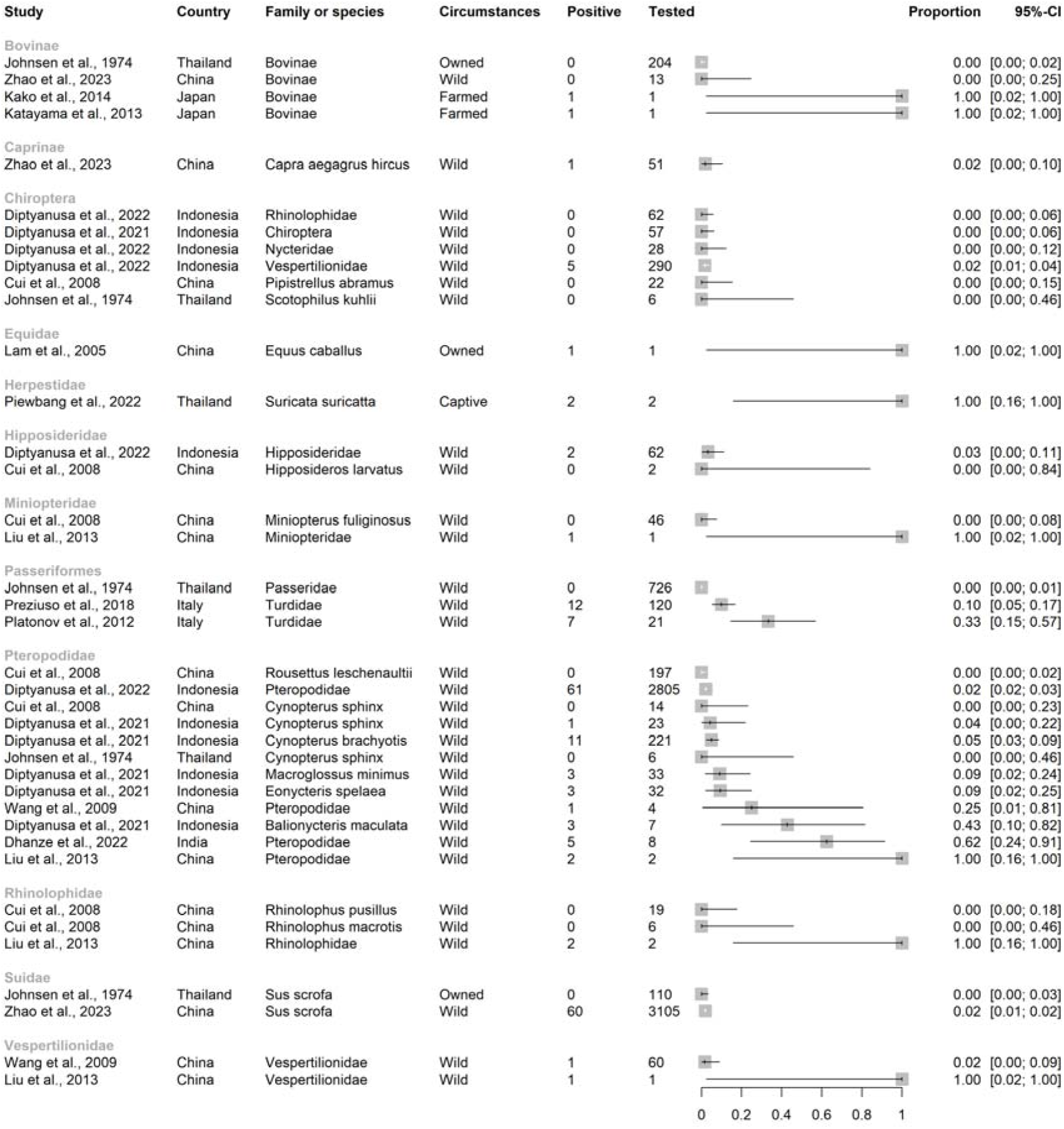
Families, subfamilies, and species with reported direct evidence of naturally occurring Japanese encephalitis virus infection in studies of vertebrate animals other than humans, ardeid birds and pigs. Suidae are included from studies in which vertebrate animals other than solely humans, ardeid birds and pigs were also tested. Horizontal lines = 95% confidence intervals.

**Figure 6.**
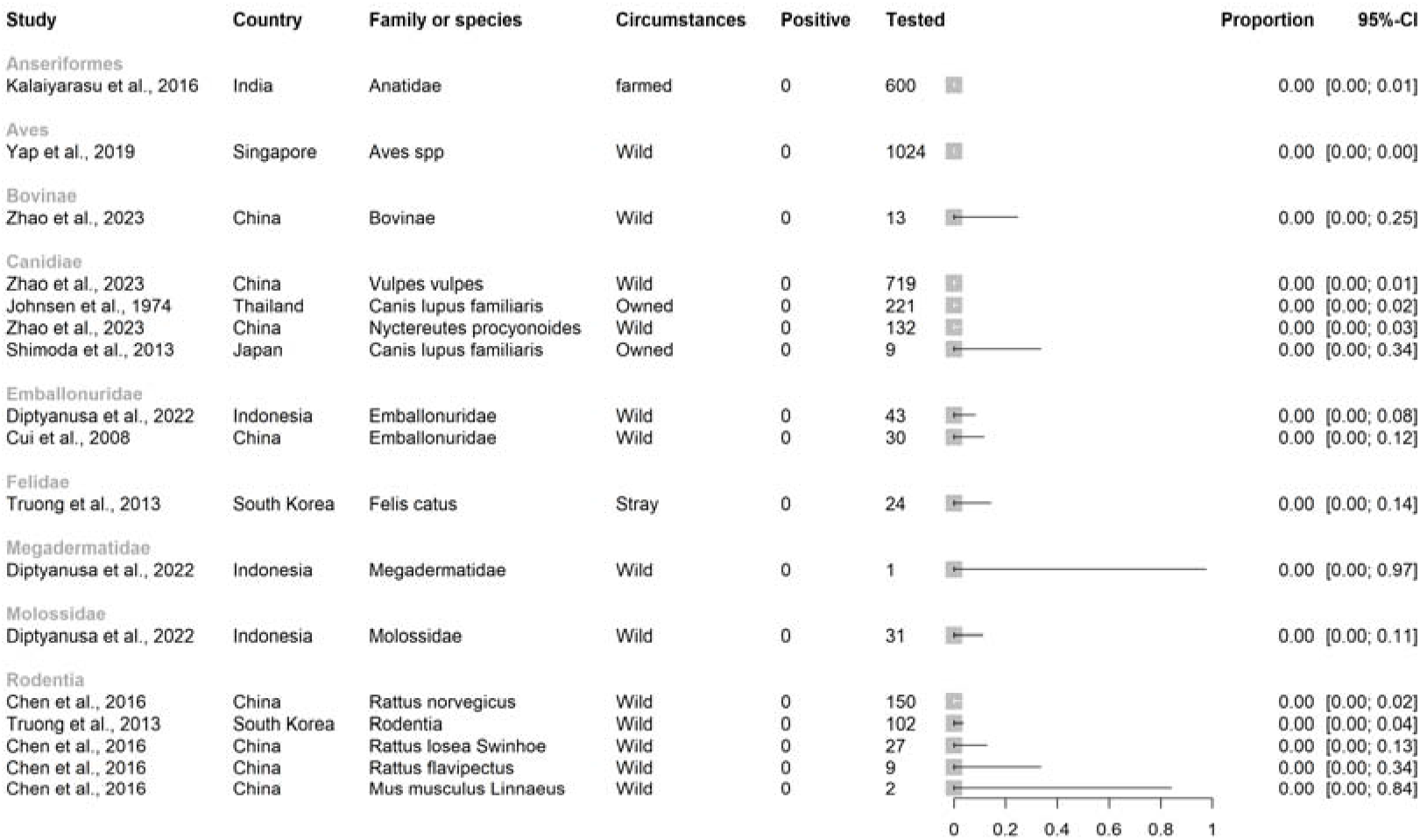
Orders and families in in studies in which evidence of JEV infection was not detected in a scoping review of Japanese encephalitis virus infection in vertebrate animals other than humans, pigs and ardeid birds. Horizontal lines = 95% confidence intervals.

Japanese encephalitis virus was also detected in other mammals including farmed Caprinae and Suidae in China (each with prevalence 2% in a survey which also included fox [*Vulpes vulpes*], racoon dog [N*yctereutes procyonoides*] and yak [*Bos grunniens*] in which no direct evidence of JEV infection was detected [57]), a single Thoroughbred racehorse in Hong Kong (*Equus caballus* [42]), cattle in Japan [43, 45], and two pet meerkats in Thailand (*S. suricatta* [46]). The racehorse had shown signs of severe neurological disease and viral RNA consistent with both Genotype I and II JEV strains was detected in spinal cord tissue by RT-PCR targeting the NS5 and E genes. Along with a rising JEV antibody titre, it was considered that the clinical signs had been consistent with JEV infection. The cattle in Japan were two Japanese black cattle (*B. taurus*; 141-day-old calf in Miyazaki Prefecture in 2009 and a 114-month-old beef cow in Aichi Prefecture in 2010) that had both shown neurological signs; sequenced cerebral virus isolates were consistent with JEV Group 1. The meerkats had died after developing signs of neurological disease. Brain samples were positive for JEV by immunohistochemistry and yielded JEV partial sequences that aligned with JEV strain Beijing-015, and lung samples from one meerkat also had inclusion bodies consistent with JEV infection.

Lastly, JEV was detected in samples from birds of the order Passeriformes and family Turdidae (thrushes) that were either found dead in mortality events in Tuscany, Italy 1997-2000 or were healthy birds hunted during the same period [40, 41]. In birds found dead and which were positive for JEV by IHC, JEV sequences were detected that were consistent with genotype III (RT-PCR targeting NS5 and E genes) [41]. JEV was detected in bone marrow by IHC in 10% of healthy birds (none found in kidney, spleen, liver, lung, brain, intestine), and of these, four yielded JEV sequences also most consistent with genotype III (RT-PCR targeting NS5 and E genes) [41].

#### 3.4.2. Indirect evidence of JEV

In the studies identified, 33,505 mammalian, 8738 avian, and 158 reptilian individuals were tested for indirect evidence (antibodies) of JEV infection (Figure 7). Fish and amphibia were not tested.

**Figure 7.**
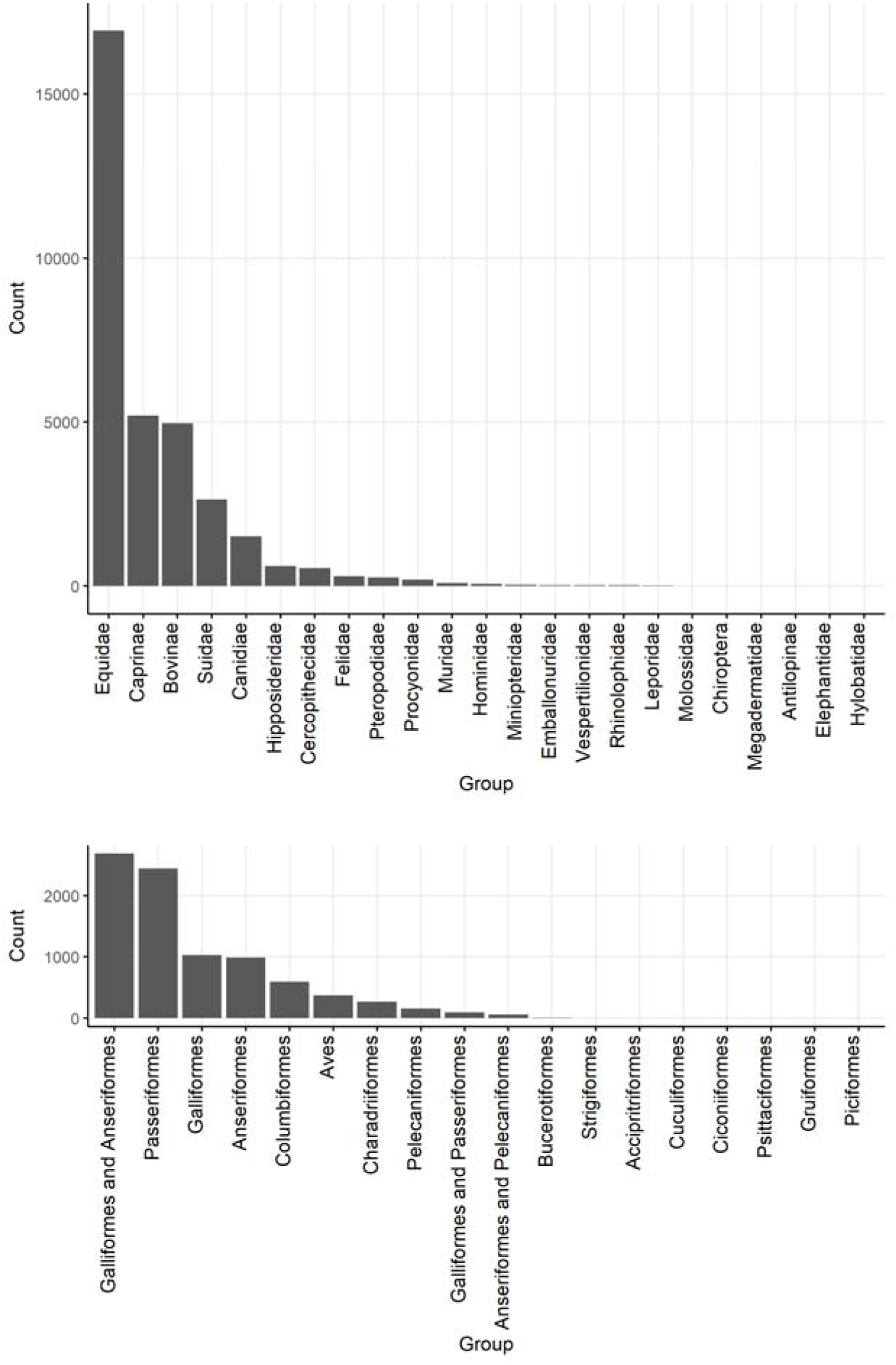
Barplots of the number of mammalian and avian individuals tested for JEV antibody in studies identified in a scoping review of direct and indirect evidence of naturally occurring Japanese encephalitis virus infection in vertebrate animals other than humans, ardeid birds and pigs.

##### 3.4.1.1 Mammalia

Of the mammalian individuals tested (Figure 7), 16,931 were horses (*E. caballus*), 5197 were Caprinae (2819 sheep [*Ovis aries*], 1 mouflon [*O. gmelina*], 2377 goats [*Capra aegagrus hircus*]), 2996 were Bovinae (at least 703 buffalo [*Bubalis bubalis*], and other cattle [*B. taurus*], or unspecified), 2633 were Suidae (mainly *Sus scrofa,* as well as 13 bearded pigs [*S. barbatus*], and four babirusa [*Babyrousa quadricornua*]), and 1516 were Canidae (>100 racoon dogs [*N. procyonoides*], one African wild dog [*Lycaon pictus*], and 1496 domestic dogs [*Canis lupus familiaris*]). Fewer than 1000 individuals were tested in each of the other families, although together, 1021 bats (order Chiroptera) were tested from the following families: Emballonuridae, Hipposideridae, Megadermatidae, Miniopteridae, Molossidae, Pteropodidae, Rhinolophidae, and Vespertilionidae.

Seroprevalence point estimates in studies ranged from 0—100% (inclusive) in subfamilies Bovinae and Caprinae and family Suidae, 0—90% in Equidae, and 0—78% in Canidae, (Figure 8).

**Figure 8.**
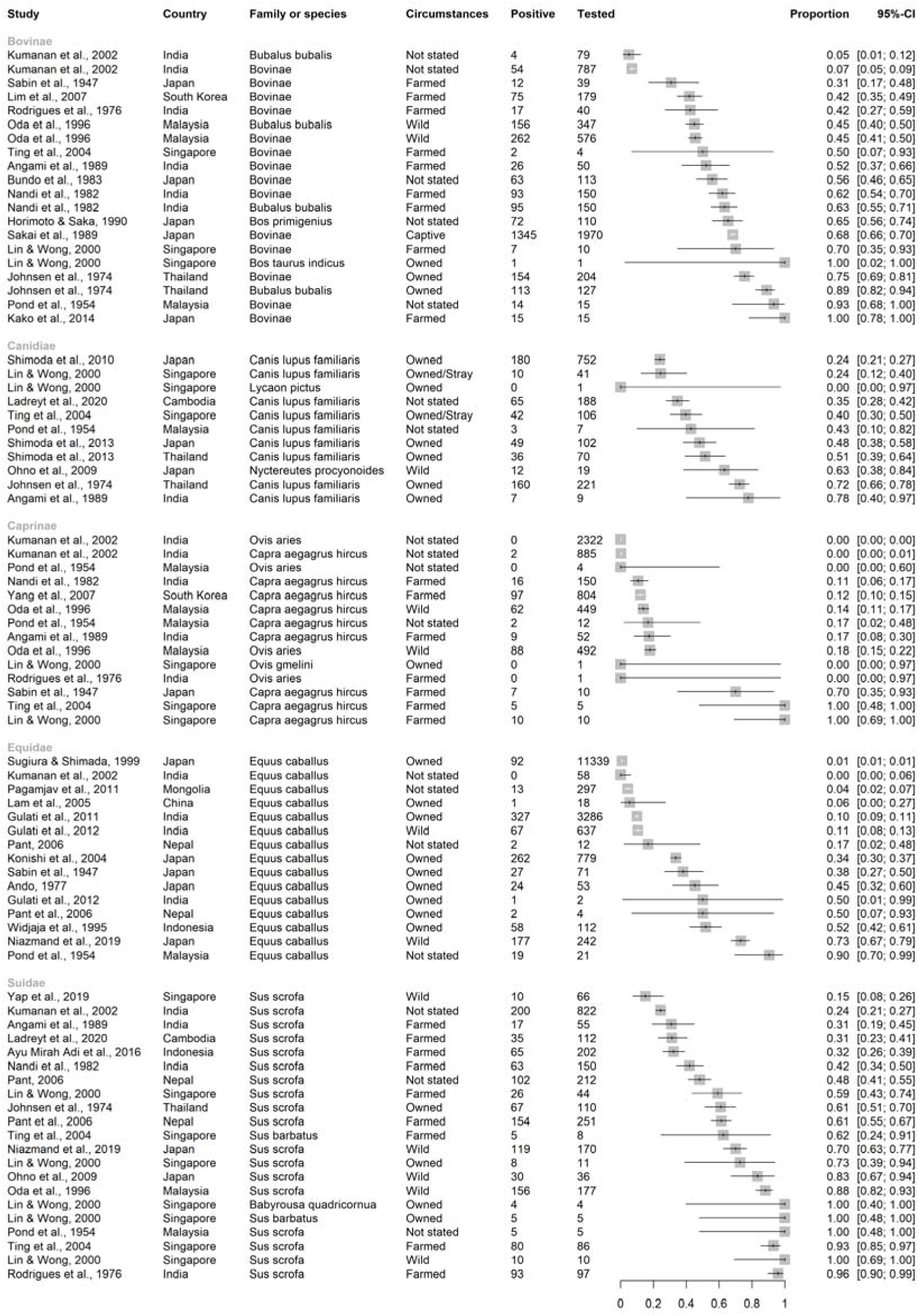
Reported seroprevalence of JEV antibodies in Bovinae, Canidae, Caprinae, Equidae, and Suidae in a scoping review of direct and indirect evidence of naturally occurring Japanese encephalitis virus infection in vertebrate animals other than humans, ardeid birds and pigs. Suidae are included only from studies in which vertebrate animals other than humans, ardeid birds or pigs were also tested. Horizontal lines = 95% confidence intervals.

Seroprevalence point estimates in studies ranged less widely in other groups of mammals (Figure S2). In studies of bats, seroprevalence ranged up to 10% in Emballonuridae, 15% in Hipposideridae (*Hipoposideros bicolor*), 11% in Miniopteridae (*Miniopterus fuliginosus*), and 18% in Pteropodidae (*Rousettus leschenaultia*) [56, 58]. Antibodies to JEV were not detected in studies of bat families Hylobatidae, Megadermatidae, Molossidae, and Rhinolophidae, and Vespertilionidae [56, 58–60]. In studies of primates, seroprevalence ranged up to 30% in Cercopithecidae (Japanese macaque, *Macaca fuscata*), and 50% in Hominidae (Bornean orangutan, *Pongo pygmaeus*) [59, 61]. In Felidae, Leporidae, Procyonidae, and Rodentia, studies detected seroprevalence up to 1%, 33%, 53% and 52%, respectively [37, 47, 62, 63]. In Singapore, antibodies to JEV were not detected in a Springbok (*Antidorcas marsupialis*; family Antilopinae), but archived serum from an *Elephas maximus* (family Elephantidae) in Singapore was positive by HIA [59].

##### 3.4.1.2 Aves

Birds were often grouped in studies so that separation of avian families by species was not possible (Figure 7). Over 1000 individuals were tested in orders Galliformes (land fowl), Passeriformes (perching birds, including songbirds), and Anseriformes (waterfowl). At least 150 ardeid birds were tested for JEV antibody alongside other species in eligible records (Pelecaniformes, including little egret [*Egretta garzetta*] and cattle egret [*Bubulcus ibis*]). The widest range of JEV seroprevalence was in studies of Anseriformes (0—86%), followed by Galliformes (0—60%), then Passeriformes (0—50%) (Figure 9). Of the Passeriformes, families in which JEV antibodies were detected were Estrilididae in Hawai’i, USA (<0.01% [64]), Passeridae in Thailand and Japan (<0.01% and 24%, respectively [65, 66]), Corvidae in Singapore and India (0.02% and 25%, respectively [67, 68], and Vangidae in India [69]. Seroprevalence in Pelecaniformes ranged from 13—22% in Ardeidae in India [68, 69]. In the study in Hawai’i, which is outside the known geographic distribution of JEV circulation, neutralising antibody consistent with JEV seropositivity was also detected in two Columbiformes (pigeons and doves; from a total of 1835 Passeriformes and Columbiformes).

**Figure 9.**
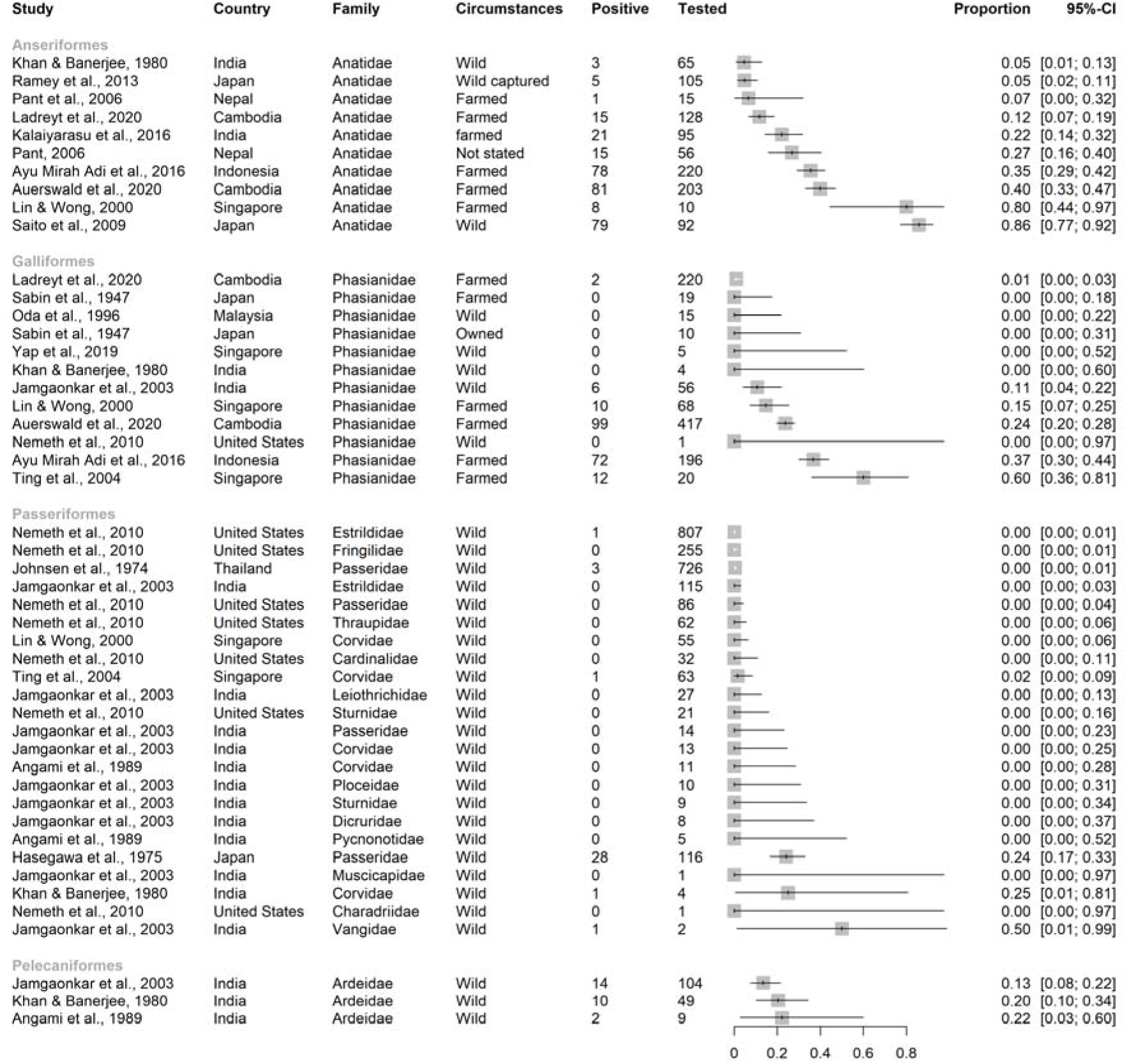
Reported seroprevalence of JEV antibodies of Anseriformes, Galliformes, Passeriformes, and Pelecaniformes in a scoping review of direct and indirect evidence of Japanese encephalitis virus infection in vertebrate animals other than humans, pigs and ardeid birds. Ardeidae (Pelecaniformes) are included from studies in which vertebrate animals other than ardeids, pigs and humans were also tested. Horizontal lines = 95% confidence intervals.

Low seroprevalence was detected in Charadriiformes (shorebirds; up to 8-10% in families Chionidae, Charadriidae, and Scolopacidae in India and Singapore [68, 70]), and Columbiformes in Hawaii, USA and India (up to 12% [39, 71]) (Figure S3). Antibody to JEV was not detected in Accipitriformes, Bucerotiformes, Ciconiiformes, Cuculiformes, Gruiformes, Piciformes, Psittaciformes, and Strigiformes in India [69].

##### 3.4.1.3 Reptilia

Eighty-two Squamata (snakes) were tested for JEV antibody. Antibodies were not detected in two reticulated pythons (*Malayopython reticulatus*) in Singapore [59]. In Kwantung Province, China, antibodies were detected in 68 and 63 Indian cobras (*Naja naja*, N = 80; seroprevalence up to 85%) by HIA and NT, respectively (80% agreement, Cohen’s kappa 0.35, which is classed as “fair” agreement), with seven cobras negative by both tests [72]. The cobras were adult females, obtained monthly in batches of 5—10 from a dealer in Hong Kong from October 1971 to September 1972. All were sourced within the previous 2 days from unknown locations in Kwantung Province. Seroprevalence and mean HIA and NT titres in cobras collected in the spring and summer months were significantly greater than those collected in the autumn and winter months.

Seventy-six Testudines (turtles) were tested for JEV antibody. Antibodies were not detected in one red-eared terrapin (*Trachemys scripta elegans*) in Singapore [59]. Seventy-five wild-caught soft-shelled turtles (*Trionyx sinensis*) from Wuchow, Kwangsi Province, China were tested for JEV antibody and 45 and 58 were positive by HIA and NT tests, respectively [73] (58% agreement, Cohen’s kappa 0.07, classed as “none to slight” agreement). Unlike the cobras from the same region, seroprevalence and titres were similar between seasons.

## 4 DISCUSSION

In this scoping review of naturally occurring JEV in vertebrates other than humans, ardeid birds and pigs, direct evidence of JEV infection through detection of virus, viral antigen or viral RNA, was found in Chiroptera (Pteropodidae, Vespertilionidae, Rhinolophidae, Miniopteridae, Hipposideridae), Passeriformes (thrushes; family Turdidae), Bovidae (cattle [*B. taurus*] and a goat [*C. hircus*]), one horse (*E. caballus*), and two meerkats (*S. suricatta*). Indirect evidence of JEV infection (antibodies) was reported for several mammalian and avian orders with high apparent seroprevalence in some studies, as well as two species of reptiles (Indian cobra [*N. naja*] and soft-shelled turtles [*T. sinensis*]). A major limitation of the evidence of naturally occurring JEV infection identified in this review was lack of information on diagnostic test accuracy, especially for detection of JEV-specific antibodies because tests can cross-react with similar orthoflaviviruses [12, 74, 75]. Due to this, as well as additional sources of heterogeneity associated with study design (for example, sampling strategies) and JEV epidemiology (for example, seasonality), study estimates were not combined. Therefore, we focus on the broader patterns of the review findings below, identifying plausible alternative hosts and surveillance opportunities rather than focusing on point estimates.

Bats are known reservoirs of many viruses – for example, Hendra virus and lyssavirus – but their role in the epidemiology of orthoflaviruses, including JEV, is unknown despite research interest [76, 77]. This interest was reflected in the current review in which bats comprised >40% of all individuals sampled for direct evidence of JEV infection over an extensive spatiotemporal window (China, India, Indonesia, and Thailand from 1974 to 2022) [49–51, 54–56, 66]. Whilst ecological studies have not found evidence that bats are involved in the epidemiology of JEV, experimental investigation indicates that bats can be competent JEV hosts [78, 79]. In the context of arboviruses, competent hosts are species in which viraemia develops to a sufficiently high titre for ongoing transmission to other individuals via the vector [80, 81]. Well-known competent hosts for JEV include ardeid birds and pigs, in which a short viraemia (duration 3—5 days) starting 1—5 days post-challenge has been shown in experimental studies [82–85]. In an experimental study based in Australia, seroprevalence in ten flying foxes (*P. alecto*) was 60% following exposure to JEV-infected mosquitoes (*Cx. annulirostris*), and although viraemia was not detected in these flying foxes, ongoing JEV transmission was demonstrated to JEV-naïve mosquitoes that fed on two of the flying foxes [78]. Given this and the direct evidence of JEV infection in bats identified in the current review, it is plausible that JEV circulation could circulate in bat populations with occasional spillover to other species. Viruses that are maintained in bat populations at low titres and prevalence might have increased replication and shedding during periods of allostatic overload on the host (cumulative and increased energy needs with external stressors such as habitat degradation and fragmentation, as well as climate anomalies), as recently described for Hendra virus [86]. If JEV infection in bat populations follows a similar mechanism, spillover of JEV from bat populations to local ardeid birds – and potentially other populations such as people, pigs, other domestic or wildlife species – during periods of allostatic overload could occur [87]. Therefore, whilst not an identified driver of detected JEV outbreaks, the role of bats in the circulation of JEV in different ecological scenarios should be considered. Another mechanism by which bats might be involved in JEV persistence or apparent re-emergence is overwintering of the virus in regions in which vector mosquitoes are seasonally inactive. To conserve energy, bats within the families Vespertilionidae and Rhinolophidae exhibit daily torpor and seasonal hibernation during cold nights and winter months, coinciding with decreased vector activity [76]. Altered immune capacity during these periods is thought to allow viruses to remain blood-borne and infectious, so that re-infection of vectors can occur post-torpor or hibernation [88]. In an experimental infection of Vespertilionidae (big brown bats [*Eptesicus fuscus*] and little brown bats [*Myotis lucifigus*]) with JEV, individuals maintained viraemia for 95—108 days after being subjected to temperatures mimicking hibernation (8—24°C) [89]. Overwintering of JEV in birds or pigs is not considered possible because their viraemia is short [90]. Prolonged viraemia could also occur in reptiles; in an experimental study in Japan, JEV viraemia was detected in lizards for several weeks following infection via *Culex* mosquitoes [91]. The apparent prevalence of antibodies suggestive of naturally occurring JEV infection in reptiles was also identified in the current review [72, 73]. Therefore, overwintering of JEV in bats and reptiles is another possible feature worth consideration in the epidemiology of JEV.

The potential role of Passeriformes in JEV epidemiology is also of interest because Passeriform birds are known reservoir and amplificatory hosts for other arboviruses, including Sindbis virus, Tahyna orthobunyavirus and Batai orthobunyavirus in Europe and WNV in north America [92, 93]. In the current review, direct evidence of JEV infection was found in thrushes (Turdidae) collected from mortality events or hunting in Tuscany, Italy in the late 1990s [40, 41]. Seropositivity was reported in other Passeriform families including Scolopacidae, Estrilidae, Corvidae, and Vangidae [39, 60, 67, 69, 94]. Serosurveillance of Passeriformes could be valuable, especially as a component of JEV preparedness activities in non-endemic regions. However, accurate diagnostic tests are needed to differentiate orthoflavivirus antibodies in the known geographic range of JEV such as WNV, as well as those in non-endemic regions, such as SLEV in North America, and this has been the subject of recent reviews [95, 96]. For example, in a study in Hawai’i in which neutralising antibody consistent with JEV seropositivity was detected in two Columbiformes, previous JEV infection was considered plausible due to the presence of competent hosts, vectors, and the proximity of air travel from Asia. However, cross-reactivity with WNV and SLEV antibodies or another, unknown flavivirus, could not be ruled out.

In addition, we identified numerous species that although unlikely to play a role as competent hosts, could be used as sentinels for JEV surveillance. In the current review, high seroprevalence (if JEV specific) of companion and livestock animals (cattle, dogs, goats, horses, ducks and land fowl) suggested that they could be useful for surveillance, especially given that these animals are kept in proximity to people. We found no direct evidence of naturally occurring JEV infection in waterfowl or land fowl (Anseriformes and Galliformes), despite epidemiological indication that they could be important in India and experimentally induced viraemia in ducklings – whilst they are useful in serosurveillance programs, their inclusion in JEV spread modelling as competent hosts is not well supported [19, 97].

Although studies identified in the current review focused on evidence of JEV infection in known and ‘known unknown’ alternative hosts (e.g. horses and bats, respectively), there were some unexpected findings. In species which are considered non-competent (dead-end) hosts, only horses (rarely) and people (estimated 0-5-1% of those infected) are generally considered to show clinical signs. However, disease attributable to JE was also reported in cattle (known non-competent JEV hosts) and meerkats (order Carnivora, suborder Feliformia) [43, 45, 46]. Whilst it could be expected that occasional clinical disease could occur in cattle, detection of disease in meerkats is unexpected and highlights the importance of comprehensive clinical reasoning in neurological cases.

In considering the limitations of the current review, in addition to the challenges of accurate diagnostic tests and false positive results, the adage that absence of evidence is not evidence of absence is highly applicable [98]. Studies were only included if they were in English and in peer-reviewed publications. Therefore, we would have excluded publications in languages from most JE-affected countries, and findings that are only available in local government reports or media; for example, there is a local government webpage report of a confirmed JE case in an alpaca in South Australia in 2022 [99]. In addition, studies in which animals were tested but findings were negative are as important as positive findings but are less likely to be published at all. Finally, the geographic distribution of JE-cases will affect the research effort. For example, JE has only recently emerged as a significant human and animal health concern in Australia, which has a unique marsupial fauna. Australian marsupials are known reservoirs of arboviruses including potentially flaviviruses, and macropods and possums have been experimentally infected with JEV [100, 101]; however, naturally occurring JEV in marsupials has yet to be reported.

## 5 CONCLUSION

We conclude that several species other than humans, ardeid birds and pigs may contribute to the epidemiology of JEV. Potential competent hosts included passerine birds and bats. We hypothesise that bats in particular might be important additional hosts to consider in JEV epidemiology and we recommend that JEV surveillance be extended to bats. Given intensifying anthropogenic ecological stressors and the propensity of orthoflaviruses for emergence, this could be of increasing importance. This review also demonstrates a range of species for serosurveillance. These included known companion and livestock non-competent hosts, as well as species which might be less frequently considered in serosurveillance programs (for example, passerine birds). However, we also demonstrated a critical need for more accurate diagnostic tests to differentiate co-circulating cross-reactive flaviviruses. Nearly all studies highlighted this as a limitation of antibody testing, and this is currently a major limitation not only in diagnosis of JEV infection, but also for JEV serosurveillance.

## Acknowledgements

We thank the librarians at the University of Sydney for their assistance with this study. This study was completed in partial fulfilment of the requirements of the Doctor of Veterinary Medicine degree, The University of Sydney (ZL).

