## Supplementary material for "A scoping review of evidence of naturally occurring Japanese encephalitis infection in vertebrate animals other than humans, ardeid birds and pigs": Table 2 Figures S1-7

**Table S1** Search terms in a scoping review of direct and indirect evidence of naturally occurring Japanese encephalitis virus infection in vertebrate animals other than humans, ardeid birds and pigs..

| Latest search date | 27 January 2024 |
|---|---|
| Web of Science (all databases), https://www.webofscience.com/ | "Japanese Encephalitis" OR JEV OR JE (Title) and detection OR detected (Topic) |
| Scopus, https://www.scopus.com/ | "Japanese Encephalitis" OR jev OR je (Article title) and detection OR detected (All fields) |
| ProQuest Central, https://www.proquest.com/central | title("Japanese encephalitis" OR "JEV" OR "JE") AND (detection OR detected); Peer reviewed |
| Google Scholar, https://scholar.google.com.au/ | Exact phrase in title: "Japanese encephalitis"; date range 1935-1980 |

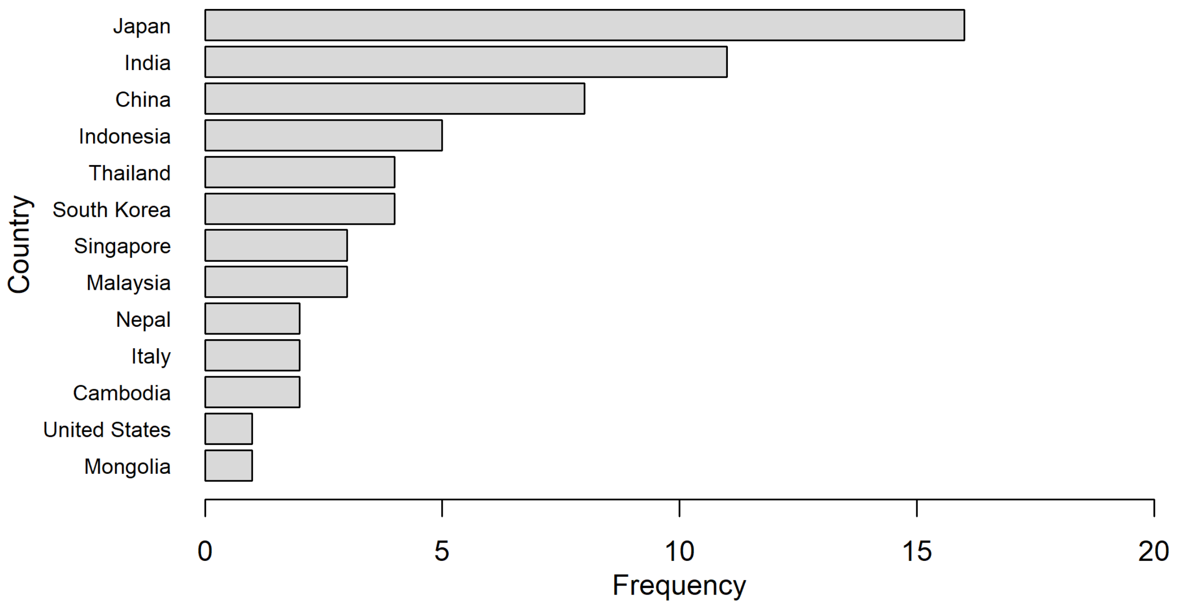


**Figure S1** Bar plot of the frequency of study location by country, in a scoping review of direct and indirect evidence of naturally occurring Japanese encephalitis virus infection in vertebrate animals other than humans, ardeid birds and pigs.

| Study | Country | Species | Circumstances | Positive | Tested | Proportion | 95%-CI |
|---|---|---|---|---|---|---|---|
| **Antilopinae** | | | | | | | |
| Lin & Wong, 2000 | Singapore | Antidorcas marsupialis | Owned | 0 | 1 | 0.00 | [0.00; 0.97] |
| **Cercopithecidae** | | | | | | | |
| Yuwono et al., 1984 | Japan | Macaca radiata | Wild | 0 | 33 | 0.00 | [0.00; 0.11] |
| Yuwono et al., 1984 | Japan | Macaca fascicularis | Wild | 18 | 358 | 0.05 | [0.03; 0.08] |
| Lin & Wong, 2000 | Singapore | Macaca fascicularis | Owned | 0 | 3 | 0.00 | [0.00; 0.71] |
| Nakgoi et al., 2014 | Thailand | Macaca nemestrina | Captive | 5 | 38 | 0.13 | [0.04; 0.28] |
| Niazmand et al., 2019 | Japan | Macaca fuscata | Wild | 22 | 75 | 0.29 | [0.19; 0.41] |
| Yuwono et al., 1984 | Japan | Macaca fuscata | Wild | 11 | 37 | 0.30 | [0.16; 0.47] |
| **Chiroptera** | | | | | | | |
| Cui et al., 2008 | China | Pipistrellus abramus | Wild | 0 | 22 | 0.00 | [0.00; 0.15] |
| Johnsen et al., 1974 | Thailand | Scotophilus kuhlii | Wild | 0 | 5 | 0.00 | [0.00; 0.52] |
| Lin & Wong, 2000 | Singapore | Chiroptera | Owned | 0 | 5 | 0.00 | [0.00; 0.52] |
| **Elephantidae** | | | | | | | |
| Lin & Wong, 2000 | Singapore | Elephas maximus | Owned | 1 | 1 | 1.00 | [0.02; 1.00] |
| **Emballonuridae** | | | | | | | |
| Cui et al., 2008 | China | Emballonuridae | Wild | 3 | 30 | 0.10 | [0.02; 0.27] |
| Johnsen et al., 1974 | Thailand | Taphozous longimanus | Wild | 0 | 2 | 0.00 | [0.00; 0.84] |
| **Felidae** | | | | | | | |
| Shimoda et al., 2010 | Japan | Felis catus | Owned | 3 | 292 | 0.01 | [0.00; 0.03] |
| Lin & Wong, 2000 | Singapore | Felis catus | Stray | 0 | 10 | 0.00 | [0.00; 0.31] |
| Lin & Wong, 2000 | Singapore | Panthera pardus | Owned | 0 | 1 | 0.00 | [0.00; 0.97] |
| Lin & Wong, 2000 | Singapore | Puma concolor | Owned | 0 | 1 | 0.00 | [0.00; 0.97] |
| **Hipposideridae** | | | | | | | |
| Banerjee et al., 1988 | India | Hipposideros pomona | Wild captured | 3 | 59 | 0.05 | [0.01; 0.14] |
| Banerjee et al., 1988 | India | Hipposideros rouxi | Wild captured | 19 | 302 | 0.06 | [0.04; 0.10] |
| Banerjee et al., 1988 | India | Hipposideros speoris | Wild captured | 12 | 167 | 0.07 | [0.04; 0.12] |
| Banerjee et al., 1988 | India | Hipposideros cineraceus | Wild captured | 2 | 22 | 0.09 | [0.01; 0.29] |
| Banerjee et al., 1988 | India | Hipposideros bicolor | Wild captured | 10 | 65 | 0.15 | [0.08; 0.26] |
| Cui et al., 2008 | China | Hipposideros larvatus | Wild | 0 | 2 | 0.00 | [0.00; 0.84] |
| **Hominidae** | | | | | | | |
| Wolfe et al., 2001 | Malaysia | Pongo pygmaeus | Semi-captive | 5 | 31 | 0.16 | [0.05; 0.34] |
| Wolfe et al., 2001 | Malaysia | Pongo pygmaeus | Wild | 15 | 40 | 0.38 | [0.23; 0.54] |
| Lin & Wong, 2000 | Singapore | Pongo pygmaeus | Owned | 1 | 2 | 0.50 | [0.01; 0.99] |
| **Hylobatidae** | | | | | | | |
| Lin & Wong, 2000 | Singapore | Symphalangus syndactylus | Owned | 0 | 1 | 0.00 | [0.00; 0.97] |
| **Leporidae** | | | | | | | |
| Kumanan et al., 2002 | India | Oryctolagus cuniculus | Not stated | 0 | 11 | 0.00 | [0.00; 0.28] |
| Sabin et al., 1947 | Japan | Oryctolagus cuniculus | Farmed | 2 | 6 | 0.33 | [0.04; 0.78] |
| **Megadermatidae** | | | | | | | |
| Banerjee et al., 1988 | India | Megaderma lyra | Wild captured | 0 | 2 | 0.00 | [0.00; 0.84] |
| **Miniopteridae** | | | | | | | |
| Cui et al., 2008 | China | Miniopterus fuliginosus | Wild | 5 | 46 | 0.11 | [0.04; 0.24] |
| **Molossidae** | | | | | | | |
| Banerjee et al., 1988 | India | Tadarida aegyptiaca | Wild captured | 0 | 9 | 0.00 | [0.00; 0.34] |
| **Procyonidae** | | | | | | | |
| Ohno et al., 2009 | Japan | Procyon lotor | Wild | 109 | 204 | 0.53 | [0.46; 0.60] |
| **Pteropodidae** | | | | | | | |
| Johnsen et al., 1974 | Thailand | Rousettus leschenaultii | Wild | 0 | 15 | 0.00 | [0.00; 0.22] |
| Cui et al., 2008 | China | Cynopterus sphinx | Wild | 0 | 14 | 0.00 | [0.00; 0.23] |
| Johnsen et al., 1974 | Thailand | Eonycteris spelaea | Wild | 0 | 14 | 0.00 | [0.00; 0.23] |
| Johnsen et al., 1974 | Thailand | Cynopterus sphinx | Wild | 2 | 18 | 0.11 | [0.01; 0.35] |
| Cui et al., 2008 | China | Rousettus leschenaultii | Wild | 35 | 197 | 0.18 | [0.13; 0.24] |
| **Rhinolophidae** | | | | | | | |
| Cui et al., 2008 | China | Rhinolophus pusillus | Wild | 0 | 19 | 0.00 | [0.00; 0.18] |
| Cui et al., 2008 | China | Rhinolophus macrotis | Wild | 0 | 6 | 0.00 | [0.00; 0.46] |
| **Rodentia** | | | | | | | |
| Chen et al., 2016 | China | Rattus flavipectus | Wild | 3 | 10 | 0.30 | [0.07; 0.65] |
| Chen et al., 2016 | China | Rattus norvegicus | Wild | 28 | 61 | 0.46 | [0.33; 0.59] |
| Chen et al., 2016 | China | Mus musculus Linnaeus | Wild | 1 | 2 | 0.50 | [0.01; 0.99] |
| Chen et al., 2016 | China | Rattus losea Swinhoe | Wild | 12 | 23 | 0.52 | [0.31; 0.73] |

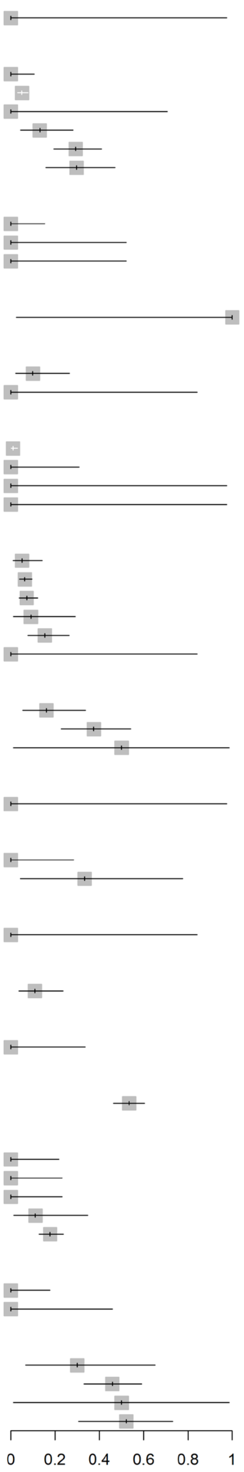


**Figure S2** Reported seroprevalence (indirect detection of JEV infection [antibody]) in studies of mammals other than Bovinae, Canidae, Caprinae, Equidae, Suidae, humans and ardeid birds. Horizontal lines = 95% confidence intervals.

| Study | Country | Family | Circumstances | Positive | Tested | Proportion | 95%-CI |
|---|---|---|---|---|---|---|---|
| **Accipritriformes** | | | | | | | |
| Jamgaonkar et al., 2003 | India | Accipitridae | Wild | 0 | 3 | 0.00 | [0.00; 0.71] |
| **Anseriformes and Pelecaniformes** | | | | | | | |
| Kumanan et al., 2002 | India | Anatidae and Ardeidae | Not stated | 6 | 64 | 0.09 | [0.04; 0.19] |
| **Aves** | | | | | | | |
| Yap et al., 2019 | Singapore | Aves spp | Wild | 12 | 371 | 0.03 | [0.02; 0.06] |
| Khan & Banerjee, 1980 | India | Aves spp | Wild | 0 | 8 | 0.00 | [0.00; 0.37] |
| **Bucerotiformes** | | | | | | | |
| Jamgaonkar et al., 2003 | India | Upupidae | Wild | 0 | 6 | 0.00 | [0.00; 0.46] |
| **Charadriiformes** | | | | | | | |
| Khan & Banerjee, 1980 | India | Chionidae | Wild | 1 | 13 | 0.08 | [0.00; 0.36] |
| Yap et al., 2019 | Singapore | Charadriidae | Wild | 6 | 69 | 0.09 | [0.03; 0.18] |
| Yap et al., 2019 | Singapore | Scolopacidae | Wild | 18 | 185 | 0.10 | [0.06; 0.15] |
| Jamgaonkar et al., 2003 | India | Charadriidae | Wild | 0 | 3 | 0.00 | [0.00; 0.71] |
| Jamgaonkar et al., 2003 | India | Burhinidae | Wild | 0 | 1 | 0.00 | [0.00; 0.97] |
| **Ciconiiformes** | | | | | | | |
| Jamgaonkar et al., 2003 | India | Ciconiidae | Wild | 0 | 2 | 0.00 | [0.00; 0.84] |
| **Columbiformes** | | | | | | | |
| Nemeth et al., 2010 | United States | Columbidae | Wild | 2 | 570 | 0.00 | [0.00; 0.01] |
| Angami et al., 1989 | India | Columbidae | Wild | 3 | 25 | 0.12 | [0.03; 0.31] |
| Khan & Banerjee, 1980 | India | Columbidae | Wild | 0 | 3 | 0.00 | [0.00; 0.71] |
| Jamgaonkar et al., 2003 | India | Columbidae | Wild | 0 | 1 | 0.00 | [0.00; 0.97] |
| **Cuculiformes** | | | | | | | |
| Jamgaonkar et al., 2003 | India | Cuculidae | Wild | 0 | 3 | 0.00 | [0.00; 0.71] |
| **Galliformes and Anseriformes** | | | | | | | |
| Yeh et al., 2012 | South Korea | Phasianidae and Anatidae | Farmed | 125 | 2681 | 0.05 | [0.04; 0.06] |
| **Galliformes and Passeriformes** | | | | | | | |
| Oda et al., 1996 | Malaysia | Phasianidae & Passeridae | Wild | 1 | 96 | 0.01 | [0.00; 0.06] |
| **Gruiformes** | | | | | | | |
| Jamgaonkar et al., 2003 | India | Rallidae | Wild | 0 | 1 | 0.00 | [0.00; 0.97] |
| **Piciformes** | | | | | | | |
| Jamgaonkar et al., 2003 | India | Megalaima haemocephala | Wild | 0 | 1 | 0.00 | [0.00; 0.97] |
| **Psittaciformes** | | | | | | | |
| Jamgaonkar et al., 2003 | India | Psittacula krameri | Wild | 0 | 2 | 0.00 | [0.00; 0.84] |
| **Strigiformes** | | | | | | | |
| Jamgaonkar et al., 2003 | India | Strigidae | Wild | 0 | 5 | 0.00 | [0.00; 0.52] |

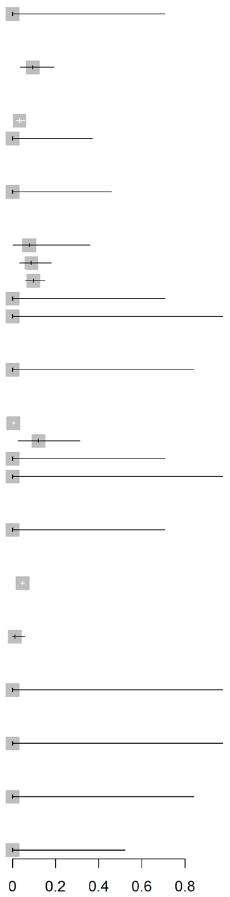


**Figure S3** Reported seroprevalence (indirect detection of JEV infection [antibody]) in bird orders in which maximum seroprevalence was <10% in a scoping review of direct and indirect evidence of naturally occurring Japanese encephalitis virus infection in vertebrate animals other than humans, ardeid birds and pigs. Ardeidae (Pelecaniformes) are included from studies in which vertebrate animals other than pigs and humans were also tested. Horizontal lines = 95% confidence intervals.

| Study | Country | Family or species | Circumstances | Positive | Tested | Proportion | 95%-CI |
|---|---|---|---|---|---|---|---|
| **China** | | | | | | | |
| Zhao et al., 2023 | China | Vulpes vulpes | Wild | 0 | 719 | 0.00 | [0.00; 0.01] |
| Cui et al., 2008 | China | Rousettus leschenaultii | Wild | 0 | 197 | 0.00 | [0.00; 0.02] |
| Chen et al., 2016 | China | Rattus norvegicus | Wild | 0 | 150 | 0.00 | [0.00; 0.02] |
| Zhao et al., 2023 | China | Nyctereutes procyonoides | Wild | 0 | 132 | 0.00 | [0.00; 0.03] |
| Cui et al., 2008 | China | Miniopterus fuliginosus | Wild | 0 | 46 | 0.00 | [0.00; 0.08] |
| Cui et al., 2008 | China | Emballonuridae | Wild | 0 | 30 | 0.00 | [0.00; 0.12] |
| Wang et al., 2009 | China | Vespertilionidae | Wild | 1 | 60 | 0.02 | [0.00; 0.09] |
| Chen et al., 2016 | China | Rattus losea Swinhoe | Wild | 0 | 27 | 0.00 | [0.00; 0.13] |
| Zhao et al., 2023 | China | Sus scrofa | Wild | 60 | 3105 | 0.02 | [0.01; 0.02] |
| Zhao et al., 2023 | China | Capra aegagrus hircus | Wild | 1 | 51 | 0.02 | [0.00; 0.10] |
| Cui et al., 2008 | China | Pipistrellus abramus | Wild | 0 | 22 | 0.00 | [0.00; 0.15] |
| Cui et al., 2008 | China | Rhinolophus pusillus | Wild | 0 | 19 | 0.00 | [0.00; 0.18] |
| Cui et al., 2008 | China | Cynopterus sphinx | Wild | 0 | 14 | 0.00 | [0.00; 0.23] |
| Zhao et al., 2023 | China | Bovinae | Wild | 0 | 13 | 0.00 | [0.00; 0.25] |
| Chen et al., 2016 | China | Rattus flavipectus | Wild | 0 | 9 | 0.00 | [0.00; 0.34] |
| Cui et al., 2008 | China | Rhinolophus macrotis | Wild | 0 | 6 | 0.00 | [0.00; 0.46] |
| Chen et al., 2016 | China | Mus musculus Linnaeus | Wild | 0 | 2 | 0.00 | [0.00; 0.84] |
| Cui et al., 2008 | China | Hipposideros larvatus | Wild | 0 | 2 | 0.00 | [0.00; 0.84] |
| Wang et al., 2009 | China | Pteropodidae | Wild | 1 | 4 | 0.25 | [0.01; 0.81] |
| Lam et al., 2005 | China | Equus caballus | Owned | 1 | 1 | 1.00 | [0.02; 1.00] |
| Liu et al., 2013 | China | Miniopteridae | Wild | 1 | 1 | 1.00 | [0.02; 1.00] |
| Liu et al., 2013 | China | Vespertilionidae | Wild | 1 | 1 | 1.00 | [0.02; 1.00] |
| Liu et al., 2013 | China | Pteropodidae | Wild | 2 | 2 | 1.00 | [0.16; 1.00] |
| Liu et al., 2013 | China | Rhinolophidae | Wild | 2 | 2 | 1.00 | [0.16; 1.00] |
| **India** | | | | | | | |
| Kalaiyarasu et al., 2016 | India | Anatidae | farmed | 0 | 600 | 0.00 | [0.00; 0.01] |
| Dhanze et al., 2022 | India | Pteropodidae | Wild | 5 | 8 | 0.62 | [0.24; 0.91] |
| **Indonesia** | | | | | | | |
| Diptyanusa et al., 2022 | Indonesia | Rhinolophidae | Wild | 0 | 62 | 0.00 | [0.00; 0.06] |
| Diptyanusa et al., 2021 | Indonesia | Chiroptera | Wild | 0 | 57 | 0.00 | [0.00; 0.06] |
| Diptyanusa et al., 2022 | Indonesia | Emballonuridae | Wild | 0 | 43 | 0.00 | [0.00; 0.08] |
| Diptyanusa et al., 2022 | Indonesia | Molossidae | Wild | 0 | 31 | 0.00 | [0.00; 0.11] |
| Diptyanusa et al., 2022 | Indonesia | Nycteridae | Wild | 0 | 28 | 0.00 | [0.00; 0.12] |
| Diptyanusa et al., 2022 | Indonesia | Vespertilionidae | Wild | 5 | 290 | 0.02 | [0.01; 0.04] |
| Diptyanusa et al., 2022 | Indonesia | Pteropodidae | Wild | 61 | 2805 | 0.02 | [0.02; 0.03] |
| Diptyanusa et al., 2022 | Indonesia | Hipposideridae | Wild | 2 | 62 | 0.03 | [0.00; 0.11] |
| Diptyanusa et al., 2021 | Indonesia | Cynopterus sphinx | Wild | 1 | 23 | 0.04 | [0.00; 0.22] |
| Diptyanusa et al., 2021 | Indonesia | Cynopterus brachyotis | Wild | 11 | 221 | 0.05 | [0.03; 0.09] |
| Diptyanusa et al., 2021 | Indonesia | Macroglossus minimus | Wild | 3 | 33 | 0.09 | [0.02; 0.24] |
| Diptyanusa et al., 2021 | Indonesia | Eonycteris spelaea | Wild | 3 | 32 | 0.09 | [0.02; 0.25] |
| Diptyanusa et al., 2022 | Indonesia | Megadermatidae | Wild | 0 | 1 | 0.00 | [0.00; 0.97] |
| Diptyanusa et al., 2021 | Indonesia | Balionycteris maculata | Wild | 3 | 7 | 0.43 | [0.10; 0.82] |
| **Italy** | | | | | | | |
| Preziuso et al., 2018 | Italy | Turdidae | Wild | 12 | 120 | 0.10 | [0.05; 0.17] |
| Platonov et al., 2012 | Italy | Turdidae | Wild | 7 | 21 | 0.33 | [0.15; 0.57] |
| **Japan** | | | | | | | |
| Shimoda et al., 2013 | Japan | Canis lupus familiaris | Owned | 0 | 9 | 0.00 | [0.00; 0.34] |
| Kako et al., 2014 | Japan | Bovinae | Farmed | 1 | 1 | 1.00 | [0.02; 1.00] |
| Katayama et al., 2013 | Japan | Bovinae | Farmed | 1 | 1 | 1.00 | [0.02; 1.00] |
| **Singapore** | | | | | | | |
| Yap et al., 2019 | Singapore | Aves spp | Wild | 0 | 1024 | 0.00 | [0.00; 0.00] |
| **South Korea** | | | | | | | |
| Truong et al., 2013 | South Korea | Rodentia | Wild | 0 | 102 | 0.00 | [0.00; 0.04] |
| Truong et al., 2013 | South Korea | Felis catus | Stray | 0 | 24 | 0.00 | [0.00; 0.14] |
| **Thailand** | | | | | | | |
| Johnsen et al., 1974 | Thailand | Passeridae | Wild | 0 | 726 | 0.00 | [0.00; 0.01] |
| Johnsen et al., 1974 | Thailand | Canis lupus familiaris | Owned | 0 | 221 | 0.00 | [0.00; 0.02] |
| Johnsen et al., 1974 | Thailand | Bovinae | Owned | 0 | 204 | 0.00 | [0.00; 0.02] |
| Johnsen et al., 1974 | Thailand | Sus scrofa | Owned | 0 | 110 | 0.00 | [0.00; 0.03] |
| Johnsen et al., 1974 | Thailand | Scotophilus kuhlii | Wild | 0 | 6 | 0.00 | [0.00; 0.46] |
| Johnsen et al., 1974 | Thailand | Cynopterus sphinx | Wild | 0 | 6 | 0.00 | [0.00; 0.46] |
| Piewbang et al., 2022 | Thailand | Suricata suricatta | Captive | 2 | 2 | 1.00 | [0.16; 1.00] |

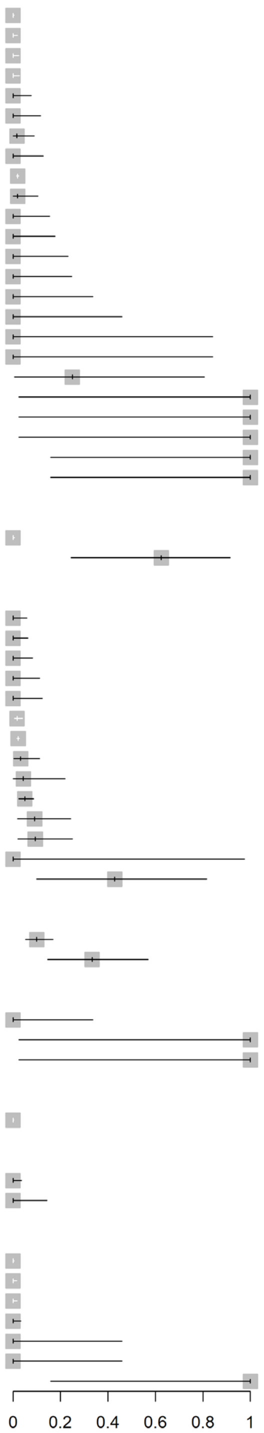


**Figure S4** Reported prevalence (direct detection of JEV infection [virus, viral antigen, or viral RNA]) stratified by country in a scoping review of direct and indirect evidence of naturally occurring Japanese encephalitis virus infection in vertebrate animals other than humans, ardeid birds and pigs. Suidae are included from studies in which vertebrate animals other than ardeid birds and humans were also tested. Horizontal lines = 95% confidence intervals.

| Study | Family | Species | Circumstances | Positive | Tested | Proportion | 95%-CI |
|---|---|---|---|---|---|---|---|
| **China** | | | | | | | |
| Cui et al., 2008 | Chiroptera | Pipistrellus abramus | Wild | 0 | 22 | 0.00 | [0.00; 0.15] |
| Cui et al., 2008 | Rhinolophidae | Rhinolophus pusillus | Wild | 0 | 19 | 0.00 | [0.00; 0.18] |
| Cui et al., 2008 | Pteropodidae | Cynopterus sphinx | Wild | 0 | 14 | 0.00 | [0.00; 0.23] |
| Lam et al., 2005 | Equidae | Equus caballus | Owned | 1 | 18 | 0.06 | [0.00; 0.27] |
| Cui et al., 2008 | Rhinolophidae | Rhinolophus macrotis | Wild | 0 | 6 | 0.00 | [0.00; 0.46] |
| Cui et al., 2008 | Emballonuridae | Emballonuridae | Wild | 3 | 30 | 0.10 | [0.02; 0.27] |
| Cui et al., 2008 | Miniopteridae | Miniopterus fuliginosus | Wild | 5 | 46 | 0.11 | [0.04; 0.24] |
| Cui et al., 2008 | Hipposideridae | Hipposideros larvatus | Wild | 0 | 2 | 0.00 | [0.00; 0.84] |
| Cui et al., 2008 | Pteropodidae | Rousettus leschenaultii | Wild | 35 | 197 | 0.18 | [0.13; 0.24] |
| Chen et al., 2016 | Rodentia | Rattus flavipectus | Wild | 3 | 10 | 0.30 | [0.07; 0.65] |
| Chen et al., 2016 | Rodentia | Rattus norvegicus | Wild | 28 | 61 | 0.46 | [0.33; 0.59] |
| Chen et al., 2016 | Rodentia | Mus musculus Linnaeus | Wild | 1 | 2 | 0.50 | [0.01; 0.99] |
| Chen et al., 2016 | Rodentia | Rattus losea Swinhoe | Wild | 12 | 23 | 0.52 | [0.31; 0.73] |
| **India** | | | | | | | |
| Kumanan et al., 2002 | Caprinae | Ovis aries | Not stated | 0 | 2322 | 0.00 | [0.00; 0.00] |
| Kumanan et al., 2002 | Caprinae | Capra aegagrus hircus | Not stated | 2 | 885 | 0.00 | [0.00; 0.01] |
| Kumanan et al., 2002 | Equidae | Equus caballus | Not stated | 0 | 58 | 0.00 | [0.00; 0.06] |
| Kumanan et al., 2002 | Leporidae | Oryctolagus cuniculus | Not stated | 0 | 11 | 0.00 | [0.00; 0.28] |
| Banerjee et al., 1988 | Molossidae | Tadarida aegyptiaca | Wild captured | 0 | 9 | 0.00 | [0.00; 0.34] |
| Kumanan et al., 2002 | Bovinae | Bubalus bubalis | Not stated | 4 | 79 | 0.05 | [0.01; 0.12] |
| Banerjee et al., 1988 | Hipposideridae | Hipposideros pomona | Wild captured | 3 | 59 | 0.05 | [0.01; 0.14] |
| Banerjee et al., 1988 | Hipposideridae | Hipposideros rouxi | Wild captured | 19 | 302 | 0.06 | [0.04; 0.10] |
| Kumanan et al., 2002 | Bovinae | Bovinae | Not stated | 54 | 787 | 0.07 | [0.05; 0.09] |
| Banerjee et al., 1988 | Hipposideridae | Hipposideros speoris | Wild captured | 12 | 167 | 0.07 | [0.04; 0.12] |
| Banerjee et al., 1988 | Hipposideridae | Hipposideros cineraceus | Wild captured | 2 | 22 | 0.09 | [0.01; 0.29] |
| Gulati et al., 2011 | Equidae | Equus caballus | Owned | 327 | 3286 | 0.10 | [0.09; 0.11] |
| Gulati et al., 2012 | Equidae | Equus caballus | Wild | 67 | 637 | 0.11 | [0.08; 0.13] |
| Nandi et al., 1982 | Caprinae | Capra aegagrus hircus | Farmed | 16 | 150 | 0.11 | [0.06; 0.17] |
| Banerjee et al., 1988 | Hipposideridae | Hipposideros bicolor | Wild captured | 10 | 65 | 0.15 | [0.08; 0.26] |
| Banerjee et al., 1988 | Megadermatidae | Megaderma lyra | Wild captured | 0 | 2 | 0.00 | [0.00; 0.84] |
| Angami et al., 1989 | Caprinae | Capra aegagrus hircus | Farmed | 9 | 52 | 0.17 | [0.08; 0.30] |
| Kumanan et al., 2002 | Suidae | Sus scrofa | Not stated | 200 | 822 | 0.24 | [0.21; 0.27] |
| Rodrigues et al., 1976 | Caprinae | Ovis aries | Farmed | 0 | 1 | 0.00 | [0.00; 0.97] |
| Angami et al., 1989 | Suidae | Sus scrofa | Farmed | 17 | 55 | 0.31 | [0.19; 0.45] |
| Nandi et al., 1982 | Suidae | Sus scrofa | Farmed | 63 | 150 | 0.42 | [0.34; 0.50] |
| Rodrigues et al., 1976 | Bovinae | Bovinae | Farmed | 17 | 40 | 0.42 | [0.27; 0.59] |
| Gulati et al., 2012 | Equidae | Equus caballus | Owned | 1 | 2 | 0.50 | [0.01; 0.99] |
| Angami et al., 1989 | Bovinae | Bovinae | Farmed | 26 | 50 | 0.52 | [0.37; 0.66] |
| Nandi et al., 1982 | Bovinae | Bovinae | Farmed | 93 | 150 | 0.62 | [0.54; 0.70] |
| Nandi et al., 1982 | Bovinae | Bubalus bubalis | Farmed | 95 | 150 | 0.63 | [0.55; 0.71] |
| Angami et al., 1989 | Canidiae | Canis lupus familiaris | Owned | 7 | 9 | 0.78 | [0.40; 0.97] |
| Rodrigues et al., 1976 | Suidae | Sus scrofa | Farmed | 93 | 97 | 0.96 | [0.90; 0.99] |
| **Japan** | | | | | | | |
| Sugiura & Shimada, 1999 | Equidae | Equus caballus | Owned | 92 | 11339 | 0.01 | [0.01; 0.01] |
| Shimoda et al., 2010 | Felidae | Felis catus | Owned | 3 | 292 | 0.01 | [0.00; 0.03] |
| Yuwono et al., 1984 | Cercopithecidae | Macaca radiata | Wild | 0 | 33 | 0.00 | [0.00; 0.11] |
| Yuwono et al., 1984 | Cercopithecidae | Macaca fascicularis | Wild | 18 | 358 | 0.05 | [0.03; 0.08] |
| Shimoda et al., 2010 | Canidiae | Canis lupus familiaris | Owned | 180 | 752 | 0.24 | [0.21; 0.27] |
| Niazmand et al., 2019 | Cercopithecidae | Macaca fuscata | Wild | 22 | 75 | 0.29 | [0.19; 0.41] |
| Yuwono et al., 1984 | Cercopithecidae | Macaca fuscata | Wild | 11 | 37 | 0.30 | [0.16; 0.47] |
| Sabin et al., 1947 | Bovinae | Bovinae | Farmed | 12 | 39 | 0.31 | [0.17; 0.48] |
| Sabin et al., 1947 | Leporidae | Oryctolagus cuniculus | Farmed | 2 | 6 | 0.33 | [0.04; 0.78] |
| Konishi et al., 2004 | Equidae | Equus caballus | Owned | 262 | 779 | 0.34 | [0.30; 0.37] |
| Sabin et al., 1947 | Equidae | Equus caballus | Owned | 27 | 71 | 0.38 | [0.27; 0.50] |
| Ando, 1977 | Equidae | Equus caballus | Owned | 24 | 53 | 0.45 | [0.32; 0.60] |
| Shimoda et al., 2013 | Canidiae | Canis lupus familiaris | Owned | 49 | 102 | 0.48 | [0.38; 0.58] |
| Ohno et al., 2009 | Procyonidae | Procyon lotor | Wild | 109 | 204 | 0.53 | [0.46; 0.60] |
| Bundo et al., 1983 | Bovinae | Bovinae | Not stated | 63 | 113 | 0.56 | [0.46; 0.65] |
| Ohno et al., 2009 | Canidiae | Nyctereutes procyonoides | Wild | 12 | 19 | 0.63 | [0.38; 0.84] |
| Horimoto & Saka, 1990 | Bovinae | Bos primigenius | Not stated | 72 | 110 | 0.65 | [0.56; 0.74] |
| Sakai et al., 1989 | Bovinae | Bovinae | Captive | 1345 | 1970 | 0.68 | [0.66; 0.70] |
| Niazmand et al., 2019 | Suidae | Sus scrofa | Wild | 119 | 170 | 0.70 | [0.63; 0.77] |
| Sabin et al., 1947 | Caprinae | Capra aegagrus hircus | Farmed | 7 | 10 | 0.70 | [0.35; 0.93] |
| Niazmand et al., 2019 | Equidae | Equus caballus | Wild | 177 | 242 | 0.73 | [0.67; 0.79] |
| Ohno et al., 2009 | Suidae | Sus scrofa | Wild | 30 | 36 | 0.83 | [0.67; 0.94] |
| Kako et al., 2014 | Bovinae | Bovinae | Farmed | 15 | 15 | 1.00 | [0.78; 1.00] |
| **Thailand** | | | | | | | |
| Johnsen et al., 1974 | Pteropodidae | Rousettus leschenaultii | Wild | 0 | 15 | 0.00 | [0.00; 0.22] |
| Johnsen et al., 1974 | Pteropodidae | Eonycteris spelaea | Wild | 0 | 14 | 0.00 | [0.00; 0.23] |
| Johnsen et al., 1974 | Chiroptera | Scotophilus kuhlii | Wild | 0 | 5 | 0.00 | [0.00; 0.52] |
| Johnsen et al., 1974 | Pteropodidae | Cynopterus sphinx | Wild | 2 | 18 | 0.11 | [0.01; 0.35] |
| Nakgoi et al., 2014 | Cercopithecidae | Macaca nemestrina | Captive | 5 | 38 | 0.13 | [0.04; 0.28] |
| Johnsen et al., 1974 | Emballonuridae | Taphozous longimanus | Wild | 0 | 2 | 0.00 | [0.00; 0.84] |
| Shimoda et al., 2013 | Canidiae | Canis lupus familiaris | Owned | 36 | 70 | 0.51 | [0.39; 0.64] |
| Johnsen et al., 1974 | Suidae | Sus scrofa | Owned | 67 | 110 | 0.61 | [0.51; 0.70] |
| Johnsen et al., 1974 | Canidiae | Canis lupus familiaris | Owned | 160 | 221 | 0.72 | [0.66; 0.78] |
| Johnsen et al., 1974 | Bovinae | Bovinae | Owned | 154 | 204 | 0.75 | [0.69; 0.81] |
| Johnsen et al., 1974 | Bovinae | Bubalus bubalis | Owned | 113 | 127 | 0.89 | [0.82; 0.94] |

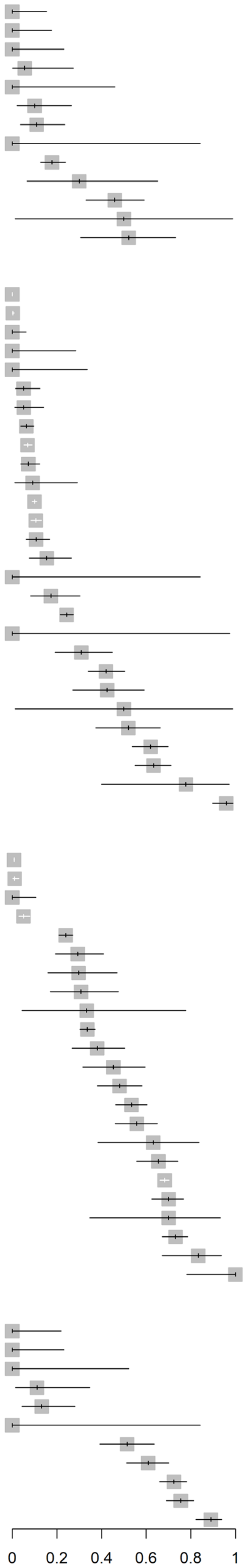


**Figure S5** Reported seroprevalence (indirect detection of JEV infection [antibody]) of mammals in China, India, Japan, and Thailand, in a scoping review of direct and indirect evidence of naturally occurring Japanese encephalitis virus infection in vertebrate animals other than humans, ardeid birds and pigs. Suidae are included from studies in which vertebrate animals other than ardeid birds and humans were also tested. Horizontal lines = 95% confidence intervals.

| Study | Family | Species | Circumstances | Positive | Tested | Proportion | 95%-CI |
|---|---|---|---|---|---|---|---|
| **Cambodia** | | | | | | | |
| Ladreyt et al., 2020 | Suidae | Sus scrofa | Farmed | 35 | 112 | 0.31 | [0.23; 0.41] |
| Ladreyt et al., 2020 | Canidiae | Canis lupus familiaris | Not stated | 65 | 188 | 0.35 | [0.28; 0.42] |
| **Indonesia** | | | | | | | |
| Ayu Mirah Adi et al., 2016 | Suidae | Sus scrofa | Farmed | 65 | 202 | 0.32 | [0.26; 0.39] |
| Widjaja et al., 1995 | Equidae | Equus caballus | Owned | 58 | 112 | 0.52 | [0.42; 0.61] |
| **Malaysia** | | | | | | | |
| Pond et al., 1954 | Caprinae | Ovis aries | Not stated | 0 | 4 | 0.00 | [0.00; 0.60] |
| Oda et al., 1996 | Caprinae | Capra aegagrus hircus | Wild | 62 | 449 | 0.14 | [0.11; 0.17] |
| Wolfe et al., 2001 | Hominidae | Pongo pygmaeus | Semi-captive | 5 | 31 | 0.16 | [0.05; 0.34] |
| Pond et al., 1954 | Caprinae | Capra aegagrus hircus | Not stated | 2 | 12 | 0.17 | [0.02; 0.48] |
| Oda et al., 1996 | Caprinae | Ovis aries | Wild | 88 | 492 | 0.18 | [0.15; 0.22] |
| Wolfe et al., 2001 | Hominidae | Pongo pygmaeus | Wild | 15 | 40 | 0.38 | [0.23; 0.54] |
| Pond et al., 1954 | Canidiae | Canis lupus familiaris | Not stated | 3 | 7 | 0.43 | [0.10; 0.82] |
| Oda et al., 1996 | Bovinae | Bubalus bubalis | Wild | 156 | 347 | 0.45 | [0.40; 0.50] |
| Oda et al., 1996 | Bovinae | Bovinae | Wild | 262 | 576 | 0.45 | [0.41; 0.50] |
| Oda et al., 1996 | Suidae | Sus scrofa | Wild | 156 | 177 | 0.88 | [0.82; 0.93] |
| Pond et al., 1954 | Equidae | Equus caballus | Not stated | 19 | 21 | 0.90 | [0.70; 0.99] |
| Pond et al., 1954 | Suidae | Sus scrofa | Not stated | 5 | 5 | 1.00 | [0.48; 1.00] |
| Pond et al., 1954 | Bovinae | Bovinae | Not stated | 14 | 15 | 0.93 | [0.68; 1.00] |
| **Mongolia** | | | | | | | |
| Pagamjav et al., 2011 | Equidae | Equus caballus | Not stated | 13 | 297 | 0.04 | [0.02; 0.07] |
| **Nepal** | | | | | | | |
| Pant, 2006 | Equidae | Equus caballus | Not stated | 2 | 12 | 0.17 | [0.02; 0.48] |
| Pant, 2006 | Suidae | Sus scrofa | Not stated | 102 | 212 | 0.48 | [0.41; 0.55] |
| Pant et al., 2006 | Equidae | Equus caballus | Farmed | 2 | 4 | 0.50 | [0.07; 0.93] |
| Pant et al., 2006 | Suidae | Sus scrofa | Farmed | 154 | 251 | 0.61 | [0.55; 0.67] |
| **Singapore** | | | | | | | |
| Lin & Wong, 2000 | Felidae | Felis catus | Stray | 0 | 10 | 0.00 | [0.00; 0.31] |
| Lin & Wong, 2000 | Chiroptera | Chiroptera | Owned | 0 | 5 | 0.00 | [0.00; 0.52] |
| Lin & Wong, 2000 | Cercopithecidae | Macaca fascicularis | Owned | 0 | 3 | 0.00 | [0.00; 0.71] |
| Yap et al., 2019 | Suidae | Sus scrofa | Wild | 10 | 66 | 0.15 | [0.08; 0.26] |
| Lin & Wong, 2000 | Canidiae | Canis lupus familiaris | Owned/Stray | 10 | 41 | 0.24 | [0.12; 0.40] |
| Lin & Wong, 2000 | Antilopinae | Antidorcas marsupialis | Owned | 0 | 1 | 0.00 | [0.00; 0.97] |
| Lin & Wong, 2000 | Canidiae | Lycaon pictus | Owned | 0 | 1 | 0.00 | [0.00; 0.97] |
| Lin & Wong, 2000 | Caprinae | Ovis gmelini | Owned | 0 | 1 | 0.00 | [0.00; 0.97] |
| Lin & Wong, 2000 | Felidae | Panthera pardus | Owned | 0 | 1 | 0.00 | [0.00; 0.97] |
| Lin & Wong, 2000 | Felidae | Puma concolor | Owned | 0 | 1 | 0.00 | [0.00; 0.97] |
| Lin & Wong, 2000 | Hylobatidae | Symphalangus syndactylus | Owned | 0 | 1 | 0.00 | [0.00; 0.97] |
| Ting et al., 2004 | Canidiae | Canis lupus familiaris | Owned/Stray | 42 | 106 | 0.40 | [0.30; 0.50] |
| Lin & Wong, 2000 | Hominidae | Pongo pygmaeus | Owned | 1 | 2 | 0.50 | [0.01; 0.99] |
| Ting et al., 2004 | Bovinae | Bovinae | Farmed | 2 | 4 | 0.50 | [0.07; 0.93] |
| Lin & Wong, 2000 | Suidae | Sus scrofa | Farmed | 26 | 44 | 0.59 | [0.43; 0.74] |
| Ting et al., 2004 | Suidae | Sus barbatus | Farmed | 5 | 8 | 0.62 | [0.24; 0.91] |
| Lin & Wong, 2000 | Bovinae | Bovinae | Farmed | 7 | 10 | 0.70 | [0.35; 0.93] |
| Lin & Wong, 2000 | Suidae | Sus scrofa | Owned | 8 | 11 | 0.73 | [0.39; 0.94] |
| Lin & Wong, 2000 | Bovinae | Bos taurus indicus | Owned | 1 | 1 | 1.00 | [0.02; 1.00] |
| Lin & Wong, 2000 | Elephantidae | Elephas maximus | Owned | 1 | 1 | 1.00 | [0.02; 1.00] |
| Lin & Wong, 2000 | Suidae | Babyrousa quadricornua | Owned | 4 | 4 | 1.00 | [0.40; 1.00] |
| Lin & Wong, 2000 | Suidae | Sus barbatus | Owned | 5 | 5 | 1.00 | [0.48; 1.00] |
| Ting et al., 2004 | Caprinae | Capra aegagrus hircus | Farmed | 5 | 5 | 1.00 | [0.48; 1.00] |
| Ting et al., 2004 | Suidae | Sus scrofa | Farmed | 80 | 86 | 0.93 | [0.85; 0.97] |
| Lin & Wong, 2000 | Caprinae | Capra aegagrus hircus | Farmed | 10 | 10 | 1.00 | [0.69; 1.00] |
| Lin & Wong, 2000 | Suidae | Sus scrofa | Wild | 10 | 10 | 1.00 | [0.69; 1.00] |
| **South Korea** | | | | | | | |
| Yang et al., 2007 | Caprinae | Capra aegagrus hircus | Farmed | 97 | 804 | 0.12 | [0.10; 0.15] |
| Lim et al., 2007 | Bovinae | Bovinae | Farmed | 75 | 179 | 0.42 | [0.35; 0.49] |
| Yeh et al., 2012 | Equidae | Equus caballus | Owned | 1468 | 2469 | 0.59 | [0.57; 0.61] |

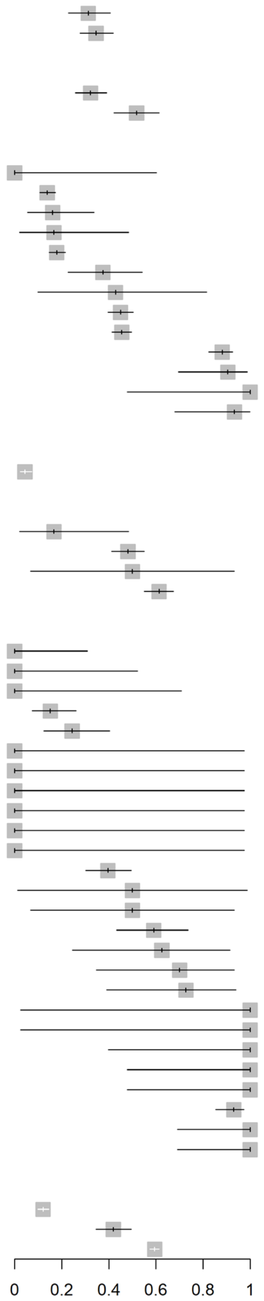


**Figure S6** Reported seroprevalence (indirect detection of JEV infection [antibody]) of mammals in Cambodia, Indonesia, Mongolia, Nepal, Singapore and South Korea in a scoping review of direct and indirect evidence of naturally occurring Japanese encephalitis virus infection in vertebrate animals other than humans, ardeid birds and pigs. Suidae are included from studies in which vertebrate animals other than ardeid birds and humans were also tested. Horizontal lines = 95% confidence intervals.

| Study by Country | Order | Family | Circumstances | Positive | Tested | Proportion | 95%-CI |
|---|---|---|---|---|---|---|---|
| **Cambodia** | | | | | | | |
| Ladreyt et al., 2020 | Galliformes | Phasianidae | Farmed | 2 | 220 | 0.01 | [0.00; 0.03] |
| Ladreyt et al., 2020 | Anseriformes | Anatidae | Farmed | 15 | 128 | 0.12 | [0.07; 0.19] |
| Auerswald et al., 2020 | Galliformes | Phasianidae | Farmed | 99 | 417 | 0.24 | [0.20; 0.28] |
| Auerswald et al., 2020 | Anseriformes | Anatidae | Farmed | 81 | 203 | 0.40 | [0.33; 0.47] |
| **India** | | | | | | | |
| Jamgaonkar et al., 2003 | Passeriformes | Estrildidae | Wild | 0 | 115 | 0.00 | [0.00; 0.03] |
| Jamgaonkar et al., 2003 | Passeriformes | Leiothrichidae | Wild | 0 | 27 | 0.00 | [0.00; 0.13] |
| Jamgaonkar et al., 2003 | Passeriformes | Passeridae | Wild | 0 | 14 | 0.00 | [0.00; 0.23] |
| Jamgaonkar et al., 2003 | Passeriformes | Corvidae | Wild | 0 | 13 | 0.00 | [0.00; 0.25] |
| Angami et al., 1989 | Passeriformes | Corvidae | Wild | 0 | 11 | 0.00 | [0.00; 0.28] |
| Jamgaonkar et al., 2003 | Passeriformes | Ploceidae | Wild | 0 | 10 | 0.00 | [0.00; 0.31] |
| Khan & Banerjee, 1980 | Anseriformes | Anatidae | Wild | 3 | 65 | 0.05 | [0.01; 0.13] |
| Jamgaonkar et al., 2003 | Passeriformes | Sturnidae | Wild | 0 | 9 | 0.00 | [0.00; 0.34] |
| Jamgaonkar et al., 2003 | Passeriformes | Dicruridae | Wild | 0 | 8 | 0.00 | [0.00; 0.37] |
| Khan & Banerjee, 1980 | Aves | Aves spp | Wild | 0 | 8 | 0.00 | [0.00; 0.37] |
| Jamgaonkar et al., 2003 | Bucerotiformes | Upupidae | Wild | 0 | 6 | 0.00 | [0.00; 0.46] |
| Khan & Banerjee, 1980 | Charadriiformes | Chionidae | Wild | 1 | 13 | 0.08 | [0.00; 0.36] |
| Angami et al., 1989 | Passeriformes | Pycnonotidae | Wild | 0 | 5 | 0.00 | [0.00; 0.52] |
| Jamgaonkar et al., 2003 | Strigiformes | Strigidae | Wild | 0 | 5 | 0.00 | [0.00; 0.52] |
| Kumanan et al., 2002 | Anseriformes and Pelecaniformes | Anatidae and Ardeidae | Not stated | 6 | 64 | 0.09 | [0.04; 0.19] |
| Khan & Banerjee, 1980 | Galliformes | Phasianidae | Wild | 0 | 4 | 0.00 | [0.00; 0.60] |
| Jamgaonkar et al., 2003 | Galliformes | Phasianidae | Wild | 6 | 56 | 0.11 | [0.04; 0.22] |
| Angami et al., 1989 | Columbiformes | Columbidae | Wild | 3 | 25 | 0.12 | [0.03; 0.31] |
| Jamgaonkar et al., 2003 | Accipritriformes | Accipitridae | Wild | 0 | 3 | 0.00 | [0.00; 0.71] |
| Jamgaonkar et al., 2003 | Charadriiformes | Charadriidae | Wild | 0 | 3 | 0.00 | [0.00; 0.71] |
| Jamgaonkar et al., 2003 | Cuculiformes | Cuculidae | Wild | 0 | 3 | 0.00 | [0.00; 0.71] |
| Khan & Banerjee, 1980 | Columbiformes | Columbidae | Wild | 0 | 3 | 0.00 | [0.00; 0.71] |
| Jamgaonkar et al., 2003 | Pelecaniformes | Ardeidae | Wild | 14 | 104 | 0.13 | [0.08; 0.22] |
| Jamgaonkar et al., 2003 | Ciconiiformes | Ciconiidae | Wild | 0 | 2 | 0.00 | [0.00; 0.84] |
| Jamgaonkar et al., 2003 | Psittaciformes | Psittacula krameri | Wild | 0 | 2 | 0.00 | [0.00; 0.84] |
| Khan & Banerjee, 1980 | Pelecaniformes | Ardeidae | Wild | 10 | 49 | 0.20 | [0.10; 0.34] |
| Kalaiyarasu et al., 2016 | Anseriformes | Anatidae | farmed | 21 | 95 | 0.22 | [0.14; 0.32] |
| Angami et al., 1989 | Pelecaniformes | Ardeidae | Wild | 2 | 9 | 0.22 | [0.03; 0.60] |
| Jamgaonkar et al., 2003 | Charadriiformes | Burhinidae | Wild | 0 | 1 | 0.00 | [0.00; 0.97] |
| Jamgaonkar et al., 2003 | Columbiformes | Columbidae | Wild | 0 | 1 | 0.00 | [0.00; 0.97] |
| Jamgaonkar et al., 2003 | Gruiformes | Rallidae | Wild | 0 | 1 | 0.00 | [0.00; 0.97] |
| Jamgaonkar et al., 2003 | Passeriformes | Muscicapidae | Wild | 0 | 1 | 0.00 | [0.00; 0.97] |
| Jamgaonkar et al., 2003 | Piciformes | Megalaima haemocephala | Wild | 0 | 1 | 0.00 | [0.00; 0.97] |
| Khan & Banerjee, 1980 | Passeriformes | Corvidae | Wild | 1 | 4 | 0.25 | [0.01; 0.81] |
| Jamgaonkar et al., 2003 | Passeriformes | Vangidae | Wild | 1 | 2 | 0.50 | [0.01; 0.99] |
| **Indonesia** | | | | | | | |
| Ayu Mirah Adi et al., 2016 | Anseriformes | Anatidae | Farmed | 78 | 220 | 0.35 | [0.29; 0.42] |
| Ayu Mirah Adi et al., 2016 | Galliformes | Phasianidae | Farmed | 72 | 196 | 0.37 | [0.30; 0.44] |
| **Japan** | | | | | | | |
| Sabin et al., 1947 | Galliformes | Phasianidae | Farmed | 0 | 19 | 0.00 | [0.00; 0.18] |
| Sabin et al., 1947 | Galliformes | Phasianidae | Owned | 0 | 10 | 0.00 | [0.00; 0.31] |
| Ramey et al., 2013 | Anseriformes | Anatidae | Wild captured | 5 | 105 | 0.05 | [0.02; 0.11] |
| Hasegawa et al., 1975 | Passeriformes | Passeridae | Wild | 28 | 116 | 0.24 | [0.17; 0.33] |
| Saito et al., 2009 | Anseriformes | Anatidae | Wild | 79 | 92 | 0.86 | [0.77; 0.92] |
| **Malaysia** | | | | | | | |
| Oda et al., 1996 | Galliformes and Passeriformes | Phasianidae & Passeridae | Wild | 1 | 96 | 0.01 | [0.00; 0.06] |
| Oda et al., 1996 | Galliformes | Phasianidae | Wild | 0 | 15 | 0.00 | [0.00; 0.22] |
| **Nepal** | | | | | | | |
| Pant et al., 2006 | Anseriformes | Anatidae | Farmed | 1 | 15 | 0.07 | [0.00; 0.32] |
| Pant, 2006 | Anseriformes | Anatidae | Not stated | 15 | 56 | 0.27 | [0.16; 0.40] |
| **Singapore** | | | | | | | |
| Lin & Wong, 2000 | Passeriformes | Corvidae | Wild | 0 | 55 | 0.00 | [0.00; 0.06] |
| Ting et al., 2004 | Passeriformes | Corvidae | Wild | 1 | 63 | 0.02 | [0.00; 0.09] |
| Yap et al., 2019 | Aves | Aves spp | Wild | 12 | 371 | 0.03 | [0.02; 0.06] |
| Yap et al., 2019 | Galliformes | Phasianidae | Wild | 0 | 5 | 0.00 | [0.00; 0.52] |
| Yap et al., 2019 | Charadriiformes | Charadriidae | Wild | 6 | 69 | 0.09 | [0.03; 0.18] |
| Yap et al., 2019 | Charadriiformes | Scolopacidae | Wild | 18 | 185 | 0.10 | [0.06; 0.15] |
| Lin & Wong, 2000 | Galliformes | Phasianidae | Farmed | 10 | 68 | 0.15 | [0.07; 0.25] |
| Ting et al., 2004 | Galliformes | Phasianidae | Farmed | 12 | 20 | 0.60 | [0.36; 0.81] |
| Lin & Wong, 2000 | Anseriformes | Anatidae | Farmed | 8 | 10 | 0.80 | [0.44; 0.97] |
| **South Korea** | | | | | | | |
| Yeh et al., 2012 | Galliformes and Anseriformes | Phasianidae and Anatidae | Farmed | 125 | 2681 | 0.05 | [0.04; 0.06] |
| **Thailand** | | | | | | | |
| Johnsen et al., 1974 | Passeriformes | Passeridae | Wild | 3 | 726 | 0.00 | [0.00; 0.01] |
| **United States** | | | | | | | |
| Nemeth et al., 2010 | Passeriformes | Estrildidae | Wild | 1 | 807 | 0.00 | [0.00; 0.01] |
| Nemeth et al., 2010 | Passeriformes | Fringilidae | Wild | 0 | 255 | 0.00 | [0.00; 0.01] |
| Nemeth et al., 2010 | Columbiformes | Columbidae | Wild | 2 | 570 | 0.00 | [0.00; 0.01] |
| Nemeth et al., 2010 | Passeriformes | Passeridae | Wild | 0 | 86 | 0.00 | [0.00; 0.04] |
| Nemeth et al., 2010 | Passeriformes | Thraupidae | Wild | 0 | 62 | 0.00 | [0.00; 0.06] |
| Nemeth et al., 2010 | Passeriformes | Cardinalidae | Wild | 0 | 32 | 0.00 | [0.00; 0.11] |
| Nemeth et al., 2010 | Passeriformes | Sturnidae | Wild | 0 | 21 | 0.00 | [0.00; 0.16] |
| Nemeth et al., 2010 | Galliformes | Phasianidae | Wild | 0 | 1 | 0.00 | [0.00; 0.97] |
| Nemeth et al., 2010 | Passeriformes | Charadriidae | Wild | 0 | 1 | 0.00 | [0.00; 0.97] |

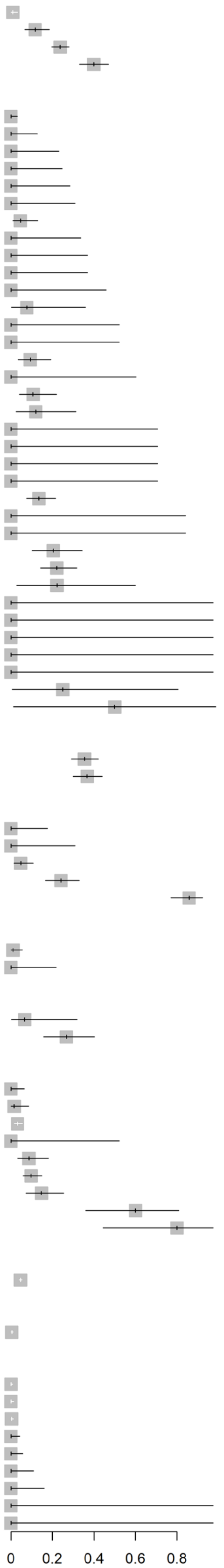


**Figure S7** Reported seroprevalence (indirect detection of JEV infection [antibody]) of birds by country in a scoping review of direct and indirect evidence of naturally occurring Japanese encephalitis virus infection in vertebrate animals other than humans, ardeid birds and pigs. Suidae are included from studies in which vertebrate animals other than ardeid birds and humans were also tested. Horizontal lines = 95% confidence intervals.
