## Supplementary material for "A scoping review of evidence of naturally occurring Japanese encephalitis infection in vertebrate animals other than humans, ardeid birds and pigs": Table S1

| Record | Earliest_YearStudy | Title | Authors | Jounal | DateStudy | Country | LocDetail | Spp | Family | Order | Class | Sick | NoTest |
|---|---|---|---|---|---|---|---|---|---|---|---|---|---|
| Ando, 1977 | 1974 | Serological epizootiology of Japanese encephalitis virus infection in horses in Japan | Ando, Y. | | 1974-1976 | Japan | Tochigi | Horse (*Equus caballus*) | Equus caballus | Perissodactyla | Mammalia | N | 53 |
| Angami et al., 1989 | 1985 | Seroepidemiological study of Japanese encephalitis in Dimapur, Nagaland | Angami, K., Chakravarty, S.K., Das, M.S., Chakraborty, M.S., Mukherjee, K.K. | The Journal of communica | Sep 1985 - Jan 1986 | India | Dimapur | Cattle | Bovinae | Artiodactyla | Mammalia | N | 50 |
| Angami et al., 1989 | 1985 | Seroepidemiological study of Japanese encephalitis in Dimapur, Nagaland | Angami, K., Chakravarty, S.K., Das, M.S., Chakraborty, M.S., Mukherjee, K.K. | The Journal of communica | Sep 1985 - Jan 1986 | India | Dimapur | Dog (Canis lupus familiaris) | Canis lupus familiaris | Carnivora | Mammalia | N | 9 |
| Angami et al., 1989 | 1985 | Seroepidemiological study of Japanese encephalitis in Dimapur, Nagaland | Angami, K., Chakravarty, S.K., Das, M.S., Chakraborty, M.S., Mukherjee, K.K. | The Journal of communica | Sep 1985 - Jan 1986 | India | Dimapur | Goat (*Capra aegagrus hircus*) | Capra aegagrus hircus | Artiodactyla | Mammalia | N | 52 |
| Angami et al., 1989 | 1985 | Seroepidemiological study of Japanese encephalitis in Dimapur, Nagaland | Angami, K., Chakravarty, S.K., Das, M.S., Chakraborty, M.S., Mukherjee, K.K. | The Journal of communica | Sep 1985 - Jan 1986 | India | Dimapur | House crow (Corvus splendens) | Corvidae | Passeriformes | Aves | N | 11 |
| Angami et al., 1989 | 1985 | Seroepidemiological study of Japanese encephalitis in Dimapur, Nagaland | Angami, K., Chakravarty, S.K., Das, M.S., Chakraborty, M.S., Mukherjee, K.K. | The Journal of communica | Sep 1985 - Jan 1986 | India | Dimapur | Little egret (*Egretta garzetta*) | Ardeidae | Pelecaniformes | Aves | N | 9 |
| Angami et al., 1989 | 1985 | Seroepidemiological study of Japanese encephalitis in Dimapur, Nagaland | Angami, K., Chakravarty, S.K., Das, M.S., Chakraborty, M.S., Mukherjee, K.K. | The Journal of communica | Sep 1985 - Jan 1986 | India | Dimapur | Pig (*Sus scrofa*) | Sus scrofa | Artiodactyla | Mammalia | N | 55 |
| Angami et al., 1989 | 1985 | Seroepidemiological study of Japanese encephalitis in Dimapur, Nagaland | Angami, K., Chakravarty, S.K., Das, M.S., Chakraborty, M.S., Mukherjee, K.K. | The Journal of communica | Sep 1985 - Jan 1986 | India | Dimapur | Pigeon (Columba livia) | Columbidae | Columbiformes | Aves | N | 14 |
| Angami et al., 1989 | 1985 | Seroepidemiological study of Japanese encephalitis in Dimapur, Nagaland | Angami, K., Chakravarty, S.K., Das, M.S., Chakraborty, M.S., Mukherjee, K.K. | The Journal of communica | Sep 1985 - Jan 1986 | India | Dimapur | Red vented bulbul (*Pycnonotus cafer*) | Pycnonotidae | Passeriformes | Aves | N | 5 |
| Angami et al., 1989 | 1985 | Seroepidemiological study of Japanese encephalitis in Dimapur, Nagaland | Angami, K., Chakravarty, S.K., Das, M.S., Chakraborty, M.S., Mukherjee, K.K. | The Journal of communica | Sep 1985 - Jan 1986 | India | Dimapur | Spotted dove (Streptopelia chinensis) | Columbidae | Columbiformes | Aves | N | 11 |
| Auerswald et al., 2020 | 2016 | Serological evidence for Japanese encephalitis and West Nile virus infections in domestic birds in Cambodia | Auerswald, H., Ruget, A-S., Ladreyt, H., In, S., Mao, S., Sorn, S., Tum, S., Duong, V., Dussart, P., Cappelle, J., & Chevalier, V. | Frontiers in veterinary scie | Mar-16 | Cambodia | Kandal, Kratie, Mondulkiri | Chicken (*Gallus gallus domesticus*) | Phasianidae | Galliformes | Aves | N | 417 |
| Auerswald et al., 2020 | 2016 | Serological evidence for Japanese encephalitis and West Nile virus infections in domestic birds in Cambodia | Auerswald, H., Ruget, A-S., Ladreyt, H., In, S., Mao, S., Sorn, S., Tum, S., Duong, V., Dussart, P., Cappelle, J., & Chevalier, V. | Frontiers in veterinary scie | Mar-16 | Cambodia | Kandal, Kratie, Mondulkiri | Duck (*Anas platyrhynchos domesticus*) | Anatidae | Anseriformes | Aves | N | 203 |
| Ayu Mirah Adi et al., 2016 | 2006 | Seroepidemiological evidence for the presence of Japanese encephalitis virus infection in ducks, chickens, and pigs, Bali-Indonesia | Ayu Mirah Adi, A., Astawa, N., Asri Damayanti, P., Kardena, I., Krisna Erawan, I., Suardana, I., Arta Putra, I., Matsumoto, Y. | Bali Medical Journal | 2006-2007 | Indonesia | Bali | Chicken (*Gallus gallus domesticus*) | Phasianidae | Galliformes | Aves | N | 196 |
| Ayu Mirah Adi et al., 2016 | 2006 | Seroepidemiological evidence for the presence of Japanese encephalitis virus infection in ducks, chickens, and pigs, Bali-Indonesia | Ayu Mirah Adi, A., Astawa, N., Asri Damayanti, P., Kardena, I., Krisna Erawan, I., Suardana, I., Arta Putra, I., Matsumoto, Y. | Bali Medical Journal | 2006-2008 | Indonesia | Bali | Duck (*Anas platyrhynchos*) | Anatidae | Anseriformes | Aves | N | 220 |
| Ayu Mirah Adi et al., 2016 | 2006 | Seroepidemiological evidence for the presence of Japanese encephalitis virus infection in ducks, chickens, and pigs, Bali-Indonesia | Ayu Mirah Adi, A., Astawa, N., Asri Damayanti, P., Kardena, I., Krisna Erawan, I., Suardana, I., Arta Putra, I., Matsumoto, Y. | Bali Medical Journal | 2006-2009 | Indonesia | Bali | Pig (*Sus scrofa*) | Sus scrofa | Artiodactyla | Mammalia | N | 202 |
| Banerjee et al., 1988 | 1983 | Antibodies against Japanese encephalitis virus in insectivorous bats from Karnataka | Banerjee, K., Bhat, H.R., Geevarghese, G., Goerge Jacob, P., & Malunjkar, A.S. | Indian journal of medical r | 1983, 1985, 1986 | India | Karanataka | Bat (*Hipposideros bicolor*) | Hipposideros bicolor | Chiroptera | Mammalia | N | 65 |
| Banerjee et al., 1988 | 1983 | Antibodies against Japanese encephalitis virus in insectivorous bats from Karnataka | Banerjee, K., Bhat, H.R., Geevarghese, G., Goerge Jacob, P., & Malunjkar, A.S. | Indian journal of medical r | 1983, 1985, 1986 | India | Karanataka | Bat (*Hipposideros cineraceus*) | Hipposideros cineraceus | Chiroptera | Mammalia | N | 22 |
| Banerjee et al., 1988 | 1983 | Antibodies against Japanese encephalitis virus in insectivorous bats from Karnataka | Banerjee, K., Bhat, H.R., Geevarghese, G., Goerge Jacob, P., & Malunjkar, A.S. | Indian journal of medical r | 1983, 1985, 1986 | India | Karanataka | Bat (*Hipposideros pomona*) | Hipposideros pomona | Chiroptera | Mammalia | N | 59 |
| Banerjee et al., 1988 | 1983 | Antibodies against Japanese encephalitis virus in insectivorous bats from Karnataka | Banerjee, K., Bhat, H.R., Geevarghese, G., Goerge Jacob, P., & Malunjkar, A.S. | Indian journal of medical r | 1983, 1985, 1986 | India | Karanataka | Bat (*Hipposideros rouxi*) | Hipposideros rouxi | Chiroptera | Mammalia | N | 302 |
| Banerjee et al., 1988 | 1983 | Antibodies against Japanese encephalitis virus in insectivorous bats from Karnataka | Banerjee, K., Bhat, H.R., Geevarghese, G., Goerge Jacob, P., & Malunjkar, A.S. | Indian journal of medical r | 1983, 1985, 1986 | India | Karanataka | Bat (*Hipposideros speoris*) | Hipposideros speoris | Chiroptera | Mammalia | N | 167 |
| Banerjee et al., 1988 | 1983 | Antibodies against Japanese encephalitis virus in insectivorous bats from Karnataka | Banerjee, K., Bhat, H.R., Geevarghese, G., Goerge Jacob, P., & Malunjkar, A.S. | Indian journal of medical r | 1983, 1985, 1986 | India | Karanataka | Bat (*Megaderma lyra*) | Megaderma lyra | Chiroptera | Mammalia | N | 2 |
| Banerjee et al., 1988 | 1983 | Antibodies against Japanese encephalitis virus in insectivorous bats from Karnataka | Banerjee, K., Bhat, H.R., Geevarghese, G., Goerge Jacob, P., & Malunjkar, A.S. | Indian journal of medical r | 1983, 1985, 1986 | India | Karanataka | Bat (*Tadarida aegyptiaca*) | Tadarida aegyptiaca | Chiroptera | Mammalia | N | 9 |
| Bundo et al., 1983 | 1981 | Antibodies against Japanese encephalitis virus in bovine sera in Nagasaki, 1981 | Bundo, K., Morita, K., & Igarashi, A. | Tropical Medicine | June-Aug 1981 | Japan | Ohseto | Bovine | Bovinae | Artiodactyla | Mammalia | N | 7 |
| Bundo et al., 1983 | 1981 | Antibodies against Japanese encephalitis virus in bovine sera in Nagasaki, 1981 | Bundo, K., Morita, K., & Igarashi, A. | Tropical Medicine | June-Aug 1981 | Japan | Azuma | Bovine | Bovinae | Artiodactyla | Mammalia | N | 17 |
| Bundo et al., 1983 | 1981 | Antibodies against Japanese encephalitis virus in bovine sera in Nagasaki, 1981 | Bundo, K., Morita, K., & Igarashi, A. | Tropical Medicine | June-Aug 1981 | Japan | Iki | Bovine | Bovinae | Artiodactyla | Mammalia | N | 20 |
| Bundo et al., 1983 | 1981 | Antibodies against Japanese encephalitis virus in bovine sera in Nagasaki, 1981 | Bundo, K., Morita, K., & Igarashi, A. | Tropical Medicine | May- Sept 1981 | Japan | Saseho | Bovine | Bovinae | Artiodactyla | Mammalia | N | 29 |
| Bundo et al., 1983 | 1981 | Antibodies against Japanese encephalitis virus in bovine sera in Nagasaki, 1981 | Bundo, K., Morita, K., & Igarashi, A. | Tropical Medicine | June-Sept 1981 | Japan | Gotoh | Bovine | Bovinae | Artiodactyla | Mammalia | N | 40 |
| Chen et al., 2016 | 2013 | Serological prevalence against Japanese encephalitis virus-serocomplex flaviviruses in commensal and field rodents in South China | Chen, S-w., Jiang, L-n., Zhong, X-s., Zheng, X-y., Ma, S-j., Xiong, Y-q., Zhou, J-h., Li, X., Ke, X-m., Zhou, W., & Chen, Q. | Vector borne and zoonotic | Nov 2013-May 2014 | China | | Brown rat (*Rattus norvegicus*) | Rattus norvegicus | Rodentia | Mammalia | N | 61 (IgG antibodies) / 25 (neutralising a |
| Chen et al., 2016 | 2013 | Serological prevalence against Japanese encephalitis virus-serocomplex flaviviruses in commensal and field rodents in South China | Chen, S-w., Jiang, L-n., Zhong, X-s., Zheng, X-y., Ma, S-j., Xiong, Y-q., Zhou, J-h., Li, X., Ke, X-m., Zhou, W., & Chen, Q. | Vector borne and zoonotic | Nov 2013-May 2014 | China | | Buff-breasted rat (*Rattus flavipectus*) | Rattus flavipectus | Rodentia | Mammalia | N | 10 (IgG antibodies) / 2 (neutralising a |
| [illegible] | | | | | | | | | | | | | |

| Record | CountIndirect | CountDirect | CountPosIndirect | CountPosDirect | Circumstances | ELISA | HIA | NT | PCR | Immunohistochemistry | Immunofluorescence | VirusSequencing | VirusIsolation | SeSp |
|---|---|---|---|---|---|---|---|---|---|---|---|---|---|---|
| Ando, 1977 | 53 | 0 | 24 | 0 | Owned | No | Yes | No | No | No | No | No | No | No |
| Angami et al., 1989 | 50 | 0 | 26 | 0 | Farmed | No | Yes | Yes | No | No | No | No | No | No |
| Angami et al., 1989 | 9 | 0 | 7 | 0 | Owned | No | Yes | Yes | No | No | No | No | No | No |
| Angami et al., 1989 | 52 | 0 | 9 | 0 | Farmed | No | Yes | Yes | No | No | No | No | No | No |
| Angami et al., 1989 | 11 | 0 | 0 | 0 | Wild | No | Yes | Yes | No | No | No | No | No | No |
| Angami et al., 1989 | 9 | 0 | 2 | 0 | Wild | No | Yes | Yes | No | No | No | No | No | No |
| Angami et al., 1989 | 55 | 0 | 17 | 0 | Farmed | No | Yes | Yes | No | No | No | No | No | No |
| Angami et al., 1989 | 14 | 0 | 3 | 0 | Wild | No | Yes | Yes | No | No | No | No | No | No |
| Angami et al., 1989 | 5 | 0 | 0 | 0 | Wild | No | Yes | Yes | No | No | No | No | No | No |
| Angami et al., 1989 | 11 | 0 | 0 | 0 | Wild | No | Yes | Yes | No | No | No | No | No | No |
| Auerswald et al., 2020 | 417 | 0 | 99 | 0 | Farmed | No | Yes | Yes | No | No | No | No | No | No |
| Auerswald et al., 2020 | 203 | 0 | 81 | 0 | Farmed | No | Yes | Yes | No | No | No | No | No | No |
| Ayu Mirah Adi et al., 2016 | 196 | 0 | 72 | 0 | Farmed | Yes | No | No | No | No | No | No | No | No |
| Ayu Mirah Adi et al., 2016 | 220 | 0 | 78 | 0 | Farmed | Yes | No | No | No | No | No | No | No | No |
| Ayu Mirah Adi et al., 2016 | 202 | 0 | 65 | 0 | Farmed | Yes | No | No | No | No | No | No | No | No |
| Banerjee et al., 1988 | 65 | 0 | 10 | 0 | Wild captured | No | No | Yes | No | No | No | No | No | No |
| Banerjee et al., 1988 | 22 | 0 | 2 | 0 | Wild captured | No | No | Yes | No | No | No | No | No | No |
| Banerjee et al., 1988 | 59 | 0 | 3 | 0 | Wild captured | No | No | Yes | No | No | No | No | No | No |
| Banerjee et al., 1988 | 302 | 0 | 19 | 0 | Wild captured | No | No | Yes | No | No | No | No | No | No |
| Banerjee et al., 1988 | 167 | 0 | 12 | 0 | Wild captured | No | No | Yes | No | No | No | No | No | No |
| Banerjee et al., 1988 | 2 | 0 | 0 | 0 | Wild captured | No | No | Yes | No | No | No | No | No | No |
| Banerjee et al., 1988 | 9 | 0 | 0 | 0 | Wild captured | No | No | Yes | No | No | No | No | No | No |
| Bundo et al., 1983 | 7 | 0 | 3 | 0 | Not stated | Yes | Yes | Yes | No | No | No | No | No | No |
| Bundo et al., 1983 | 17 | 0 | 3 | 0 | Not stated | Yes | Yes | Yes | No | No | No | No | No | No |
| Bundo et al., 1983 | 20 | 0 | 10 | 0 | Not stated | Yes | Yes | Yes | No | No | No | No | No | No |
| Bundo et al., 1983 | 29 | 0 | 19 | 0 | Not stated | Yes | Yes | Yes | No | No | No | No | No | No |
| Bundo et al., 1983 | 40 | 0 | 28 | 0 | Not stated | Yes | Yes | Yes | No | No | No | No | No | No |
| Chen et al., 2016 | 61 | 150 | 28 | 0 | Wild | Yes | No | Yes | Yes | No | No | No | No | No |
| Chen et al., 2016 | 10 | 9 | 3 | 0 | Wild | Yes | No | Yes | Yes | No | No | No | No | No |
| [illegible] | | | | | | | | | | | | | | |
